# Extremely sparse models of linkage disequilibrium in ancestrally diverse association studies

**DOI:** 10.1101/2022.09.06.506858

**Authors:** Pouria Salehi Nowbandegani, Anthony Wilder Wohns, Jenna L. Ballard, Eric S. Lander, Alex Bloemendal, Benjamin M. Neale, Luke J. O’Connor

## Abstract

Linkage disequilibrium (LD) is the correlation among nearby genetic variants. In genetic association studies, LD is often modeled using massive local correlation matrices, but this approach is slow, especially in ancestrally diverse studies. Here, we introduce *LD graphical models* (LDGMs), which are an extremely sparse and efficient representation of LD. LDGMs are derived from genome-wide genealogies; statistical relationships among alleles in the LDGM correspond to genealogical relationships among haplotypes. We publish LDGMs and ancestry specific *LDGM precision matrices* for 18 million common SNPs (MAF>1%) in five ancestry groups, validate their accuracy, and demonstrate order-of-magnitude improvements in runtime for commonly used LD matrix computations. We implement an extremely fast multi-ancestry polygenic prediction method, *BLUPx-ldgm*, which performs better than a similar method based on the reference LD correlation matrix. LDGMs will enable sophisticated methods that scale to ancestrally genetic association data across millions of variants and individuals.

## Introduction

Linkage disequilibrium (LD) is the correlation among nearby genetic variants.^1,2^ It poses a challenge in genome-wide association studies (GWAS), as disease-causing alleles reside on haplotypes with numerous tag SNPs. In applications like heritability partitioning^3,4^, polygenic risk prediction^5,6^, and fine mapping^7,8^, LD is often modeled using large local correlation matrices, with thousands of entries per SNP; however, these can be terabytes in size^8^, leading to computational bottlenecks.

The challenge is exacerbated in ancestrally diverse association studies. Diversity carries crucial scientific benefits^6,9–13^, but it also poses a methodological challenge, as LD patterns vary across ancestry groups^14^. This variation makes it even more important to model LD in applications, and it also increases the difficulty of doing so.

To model LD efficiently and accurately, a possible approach is to leverage the genealogical history that gave rise to LD in the first place. New mutations arise on haplotypes carrying existing alleles and become correlated as they increase in frequency.^1,15^ With recent breakthroughs, genome-wide genealogies of recombining organisms can be inferred from large-scale genetic datasets and recorded in *succinct tree sequences*^1–19^. Capitalizing on the limited number of common ancestral haplotypes at most loci, tree sequences provide a highly compact representation of human genetic data^18,20^. Tree sequences, and the closely related ancestral recombination graph, have enabled powerful new methods for understanding ancestral relationships^21–23^, measuring selection^19,24,25^, and analyzing complex traits^26,27^.

Here we show that LD can be modeled using extremely sparse *LD graphical models* (LDGMs) derived from inferred tree sequences. LDGMs compresses the correlation matrix in the same manner that a road map would compress a matrix of pairwise travel distances, only recording the relationship between adjacent pairs of locations or SNPs. Ancestry-specific LD patterns are encoded in an *LDGM precision matrix*, which is a sparse regularized inverse of the LD correlation matrix. We show how LDGMs address computational challenges in GWAS applications, and we apply them to analyze the transferability of polygenic risk scores across ancestries in simulations with ancestry-specific LD patterns.

## Results

### Overview of methods

Graphical models are commonly used to represent the joint distribution of several random variables. Each vertex of the graph represents a random variable, and each edge represents a conditional dependency; when two variables are not connected by an edge, it means that they are conditionally independent given the other variables. A simple example is a Markov chain, which is often used to model a time series or a sequence of states along the genome.^28^

Graphical models emerge naturally in genealogies. Suppose that four mutations, *a*, *b*, *c* and *d*, arise in a single lineage without recombination, resulting in the five haplotypes *ABCD*, *aBCD*, *abCD*, *abcD* and *abcd*. Sampling a haplotype *X* at random, the joint distribution has a Markov chain as its graphical model:

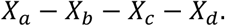

*X_a_* and *X_c_* are conditionally independent given *X_b_* because any haplotype with a *b* must have an *a*, and any haplotype with a *B* must have a *C*.

More generally, LDGMs capture the conditional dependencies that arise among SNPs when sampling from a set of ancestral haplotypes. In the Supplementary Note, we show that the

LDGM is correct, in the sense that every conditional dependency corresponds to a connected set in the LDGM, and we also show that it is minimal, in the sense that none of its edges are spurious. However, our theoretical results have two limitations related to the haplotype sampling process that we analyze. First, sampling an ancestral haplotype is not equivalent to sampling an entire genome, and in particular, population structure causes the haplotypes that are observed at different positions to be non-independent. These types of dependencies are not captured by the LDGM because they are not observed when sampling one haplotype at a time. Second, the LDGM captures all of the conditional dependencies that *may* arise in an unstructured population with certain ancestral haplotypes, but in a particular population, only some of those dependencies will actually arise (due to its particular haplotype frequencies). This distinction motivates the use of heuristics to increase the sparsity of the LDGM (in particular, an edge weight threshold and an L1 penalty; see Supplementary Note).

To construct the LDGM, we start with a tree sequence^16^ inferred using *tsinfer*^18^ and modify its edges to produce a *bricked tree sequence*. In the original tree sequence, there is a different genealogical tree at each position in the genome. The edges of each tree describe parent-child relationships between haplotypes, and the edges of the tree sequence span the entire genomic interval where a particular parent-child relationship is observed. Within that interval, the child haplotype itself may have different descendants at different positions. In the bricked tree sequence, such edges are bifurcated to form “bricks” that have the same descendants at every position. For example, in Figure 1a, there are two positions and trees, differing by a single recombination (i.e., haplotype 2 has different parents in each tree). Haplotypes 7 is the parent of haplotype 5 in both trees, but the corresponding edge is bifurcated into two bricks because haplotype 5 has different descendants in each tree (i.e., haplotype 2 is its descendent at position 1 only).

**Figure 1:**
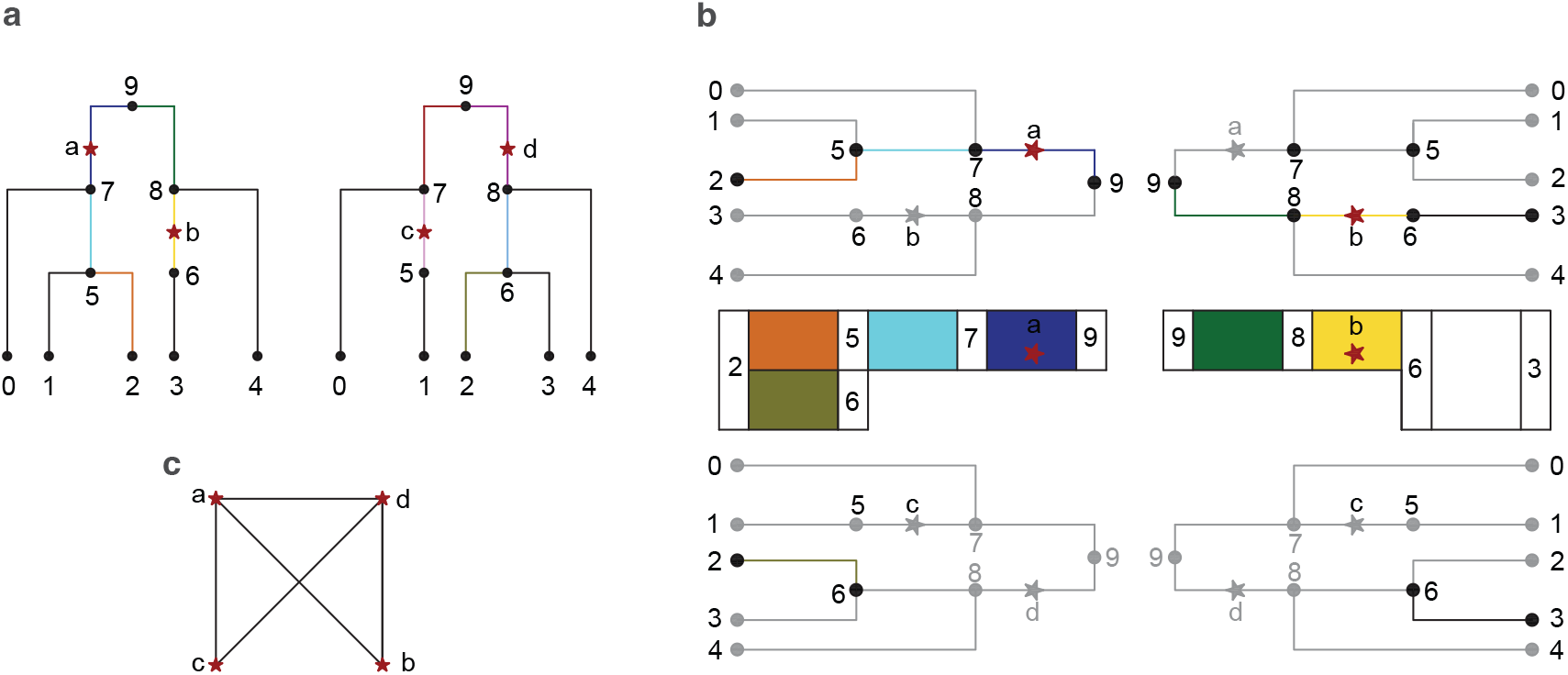
deriving LD graphical models from genome-wide genealogies. (A) First, we modify the tree sequence to produce a *bricked tree sequence*. In this example, there are two trees, differing by one recombination at haplotype 2. Some edges (colored black) span both trees. Others (denoted with colors) span only one tree; unlike in the original tree sequence, edges in the bricked tree sequence have the same descendent haplotypes in every tree that they span, and edges with this property are called “bricks.” (b) The LDGM is based on *brick diagrams*, which include all of the paths between two haplotypes via their MRCAs. The brick diagram for haplotypes 2 and 3 encodes a path with MRCA 9 and a path with MRCA 6, and it has mutations *a* and *b*. (c) The LDGM of this tree sequence has an edge between every pair of SNPs except *c* and *d*; for example, *a* and *b* are connected because they are siblings, and they comprise the symmetric difference between haplotypes 2 and 3.

The LDGM is defined using *brick diagrams*, which are generalized paths between haplotypes (Figure 1b). When there is only one genealogical tree, every two haplotypes are connected by a single path, coalescing at their most recent common ancestor (MRCA). With recombination, the haplotypes coalesce differently at different positions in the genome, and their brick diagram comprises the bricks that appear in their coalescent. Because bricks have the same descendants at every position, brick diagrams have a key property: mutations that occur on a brick diagram are carried by exactly one of the two haplotypes. This property is significant because differences between pairs of haplotypes – sets of SNPs appearing on brick diagrams – are the possible conditional dependencies of the joint distribution. The LDGM comprises a minimal set of edges guaranteeing that the SNPs on a brick diagram form a connected set in the graph (Figure 1c). We provide a formal discussion and describe our algorithm to produce the LDGM in the Supplementary Note.

Most applications of LDGMs involve *LDGM precision matrices*, which allow commonly used LD matrix operations to be performed extremely efficiently. The LDGM precision matrix is a sparse regularized inverse of the LD correlation matrix, with nonzero entries corresponding to the edges of the LDGM. Unlike the LDGM itself, the precision matrix differs across ancestry groups (because the correlation matrix is different). The precision matrix is inferred from the sample correlations corresponding to the edges of the LDGM, using a coordinate descent algorithm based on DP-GLASSO (dual-primal graphical lasso)^29^. DP-GLASSO infers a sparse precision matrix by maximizing an L1-penalized Gaussian likelihood (which is still appropriate for non-Gaussian data), but it cannot be applied to LD matrices with many thousands of SNPs. Our modified algorithm makes inference feasible by restricting to the edges of the LDGM (see Supplementary Note). The L1 penalty is used to discard edges of the LDGM that are not needed in the population being analyzed, after which the penalty is removed.

Our method involves three parameters. First, we impose a recombination frequency threshold, only bifurcating edges to form bricks if a recombined haplotype has enough descendants at the position of the recombination; we use a threshold of 1%, which is also our SNP frequency threshold. Second, we impose an edge-weight threshold, discarding edges whose weight exceeds the threshold; these weights are defined using a well-motivated heuristic, which corresponds to *r^2^* in the absence of recombination (see Supplementary Note). Third, in the precision matrix inference step, we use an L1 (lasso) penalty with a tunable coefficient to produce additional sparsity.

### Sparse and accurate LD graphical models across five ancestry groups

We analyzed high-coverage 1000 Genomes whole-genome sequencing data^30^, including 2504 individuals across five continental ancestries (African, AFR; admixed American, AMR; East Asian, EAS; European, EUR; South Asian, SAS). We included 18M SNPs with MAF>1% in any group that passed variant QC (see Methods). We inferred tree sequences using *tsinfer* (Kelleher et al., 2019; Wohns et al. 2022) (see Methods). We partitioned the genome into 1361 nearly independent LD blocks with around 13,000 SNPs per block, as previously described^32,33^, and we confirmed that these blocks were appropriate for every ancestry group (Supplementary Figure 1). One LD block containing <100 SNPs (the fewest) was removed due to QC. For each LD block, we produced one LDGM and five precision matrices (one for each ancestry group).

We quantified the accuracy and sparsity of our LDGM precision matrices across three ancestry groups in a representative LD block. As expected, LD correlation matrices vary with ancestry (Figure 2a, upper triangles). Comparing these matrices with the inverse of our LDGM precision matrices (Figure 2a, lower triangles), they are visually concordant, except for smaller nonzero entries between faraway SNPs in the LDGM precision matrix inverse. This effect is most apparent in AMR, where the correlation matrix exhibits low-rank structure due to admixture and our LDGMs are less accurate (see below). Unlike the sample correlation matrices, however, our LDGM precision matrices are extremely sparse: they have an average degree of around 20 neighbors per SNP, while most SNPs have hundreds of LD partners with *r*^2^ > 0.01 (Figure 2b).

**Figure 2:**
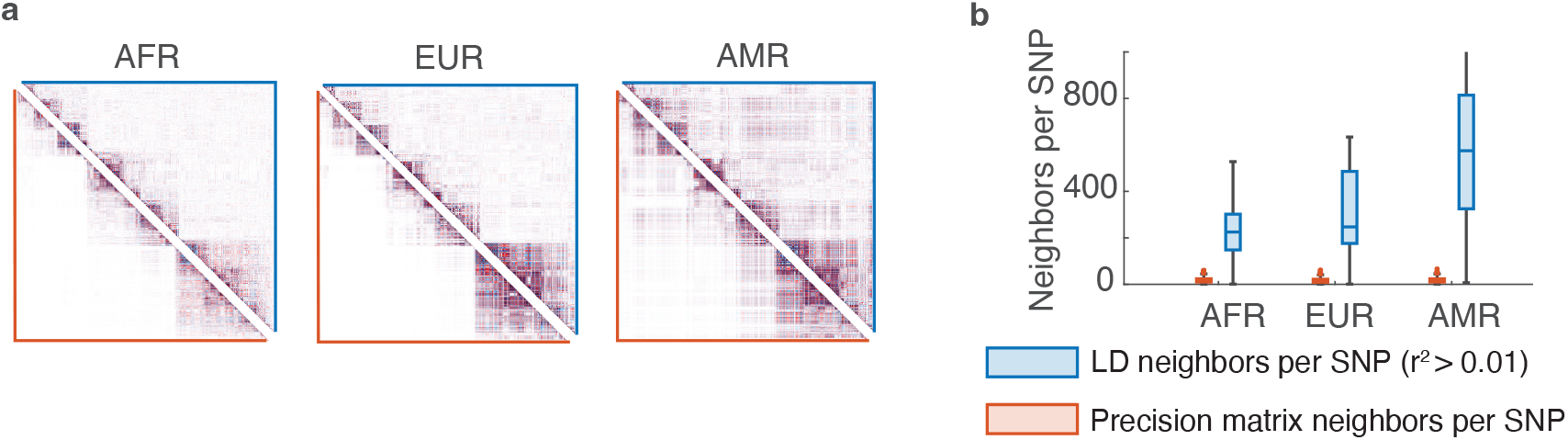
Accuracy and sparsity of the LDGM precision matrix for a representative LD block. (a) LD correlation matrices of common SNPs in three 1000 Genomes ancestry groups (AFR, AMR and EUR) (M=1,966 SNPs with MAF>1% in all ancestries). Upper triangles (blue outline) show the sample correlation matrix, and lower triangles (orange outline) show the LDGM precision matrix inverse. (b) Density of the sample correlation matrix compared with the LDGM precision matrix. We compare the number of LD partners per SNP (*r*^2^ > 0.01) in the correlation matrix with the number of nonzero entries per SNP in the precision matrix.

Similar results were obtained across the genome. Most precision matrices had an average degree of less than 20 (compared with a median block size of ~5,000) (Figure 3a), and they were highly accurate for every ancestry group except AMR, with a median mean-squared error (MSE; see Methods) of around 0.0012 (0.0023 in AMR) (Figure 3b). This number is close to the variance of the sample correlation between two uncorrelated SNPs in 500 individuals (i.e., 1/2*n* = 0.001). High accuracy was also observed using an alternative error metric (Supplementary Figure 2). The much lower accuracy observed in the admixed AMR population is expected, since LDGMs are justified mathematically under a model that does not include admixture or population structure. It suggests that LDGMs should not be currently applied to admixed datasets; however, this limitation might be addressed using local ancestry inference^34^ (see Discussion).

**Figure 3:**
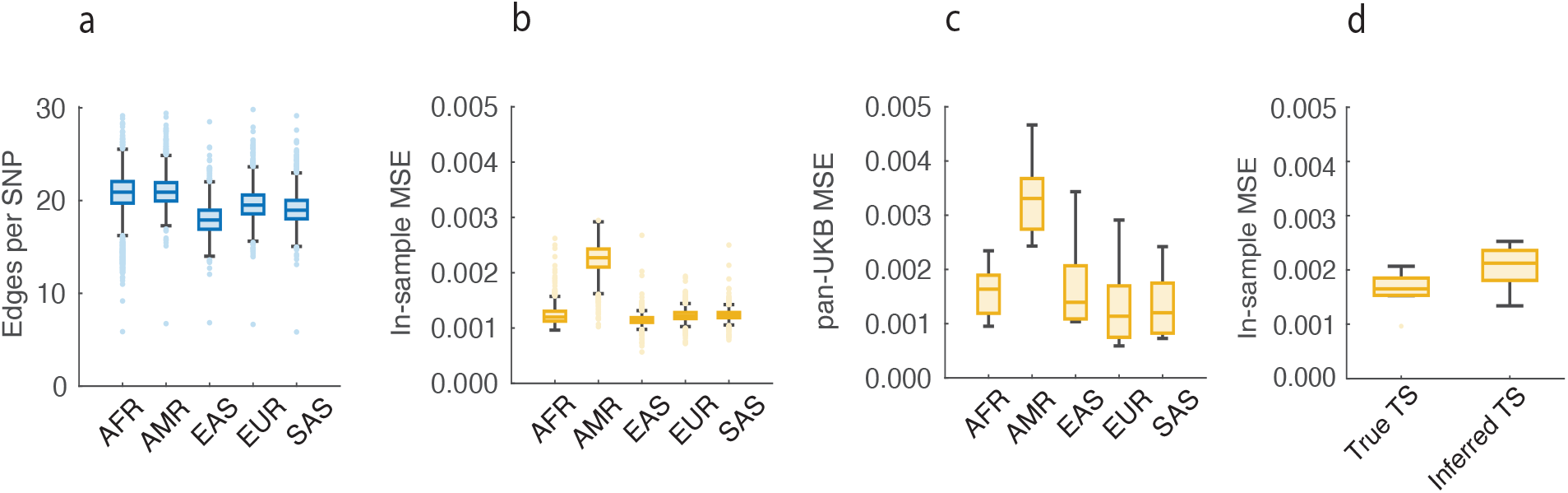
Accuracy and sparsity of LDGM precision matrices across the genome. (a) Average degree (neighbors per SNP) within each LD block across ancestries. (b) Mean squared error (MSE) of each LD block across ancestries. (c) Out-of-sample MSE of LD precision matrices on chromosome 22, comparing LDGM precision matrices derived from 1000 Genomes with ancestry-matched correlation matrices from pan-UK Biobank. (d) Accuracy and sparsity of LDGM precision matrices derived from true vs. inferred tree sequences in simulations with *msprime*. For numerical results, see Supplementary Tables 1-2.

We performed out-of-sample validation in pan-UK Biobank^35^, which includes samples from each of five subsamples based on genetic ancestry similarity (see Data Availability). Across chromosome 22, our LD precision matrices had a median out-of-sample MSE of around 0.0013 in AFR, EAS and SAS, 0.0016 in AFR, and 0.0033 in AMR (Figure 3c). These numbers support the use of 1000 genomes based LDGMs in UK Biobank, but we caution that they will not be appropriate for every dataset (see Discussion).

We investigated whether the accuracy of tree sequence inference affected the accuracy of our LDGMs. We simulated tree sequences using msprime^36^ under a neutral coalescent model with human-like parameters in regions the size of a typical LD block (see Methods). Tree sequences were inferred from the simulated genetic variation data with tsinfer^18^. Next, LDGMs and precision matrices were inferred for each simulated and inferred tree sequence. MSE was approximately 24% higher in inferred vs. simulated tree sequences (Figure 3d), indicating that imperfect tree sequence inference is a minor source of error for our LDGMs.

An existing approach to regularize the LD correlation matrix is the estimator of Wen and Stephens, which shrinks the correlation between each pair of SNPs by a factor that depends on their map distance and the sample size (but does not produce a very sparse matrix). Comparing the Wen and Stephens estimator with the inverse of our LDGM precision matrix on chromosome 22 EUR, we found that the MSE between them is very low, since both methods shrink the correlations between faraway SNPs toward zero (Supplementary Figure 3a). This similarity is encouraging, as the Wen and Stephens estimator has been shown to improve heritability estimates when it is used instead of the sample correlation matrix. Although the Wen and Stephens estimator produces sparsity when it is applied to large regions, it does not produce much sparsity within LD blocks (Supplementary Figure 3c).

A different existing approach, which does compress the local LD matrix, is to truncate its eigenvalues. Shi et al.^37,38^ used this approach to estimate heritability and genetic correlations, and it has been used by recently developed methods as well^39–41^. We compared our LDGM precision matrices with rank-*k* approximations for different values of *k* and found that LDGM precision matrices are much more accurate than small-*k* approximations (Supplementary Figure 4a-b). Unlike the Wen and Stephens estimator, low-rank approximations did not exhibit similarity with the LDGM precision matrix inverse (Supplementary Figure 4c).

We performed four secondary analyses on chromosome 22 in EUR. First, we performed twofold cross validation within 1000 Genomes, fitting our precision matrices to 50% of EUR individuals and testing in the other 50%. Median cross-validated MSE was 0.0027 (within-sample MSE: 0.0019); this number compares favorably with the MSE between the two sample correlation matrices (median: 0.0038, *P*_binomial_ < 10^-6^) (Supplementary Figure 5). The higher within-sample MSE in this analysis indicates that within-sample MSE decreases with sample size. Second, to verify that our LDGM performs better than a naïve approach, we computed precision matrices from a simple banded-diagonal LDGM (Supplementary Figure 6). Depending on band size, the resulting precision matrices either had higher MSE and similar density (median MSE: 0.003, degree: 20.1), or similar MSE and higher density (median MSE: 0.0013, degree: 38.3). We also tried an *r*^2^-threshold based LDGM, which had high MSE (median MSE: 0.0053, degree: 17.3). Third, we stratified SNPs into three allele frequency bins (0.01-0.05, 0.05-0.2, 0.2-0.5); low-frequency SNPs had slightly higher MSE (median: 0.0013, 0.0012, 0.0011 respectively) and substantially smaller average degree (median: 12.5, 19.7, 24.1) (Supplementary Figure 7). Fourth, we evaluated the performance of our LDGMs with different L1 penalties and path weight thresholds (Supplementary Figure 8). Different parameter settings produce different tradeoffs between within-sample accuracy and sparsity, and there is no choice of parameters that is obviously best.

### Computational advantages of LDGMs

Statistical methods to analyze GWAS data often utilize LD data from a reference panel together with summary association statistics.^42^ Many of these methods, especially sophisticated Bayesian algorithms, rely on slow LD matrix operations. Slow operations involving the LD correlation matrix, *R*, can be replaced with efficient operations involving the LDGM precision matrix, *P*. In Figure 4a and the Supplementary Note, we list LD matrix operations that can be replaced with the LDGM precision matrix, with their respective runtimes. For most operations, order-of-magnitude improvements are obtained. These include basic operations like matrix-by-vector multiplication and also more complex calculations involving the sampling distribution of the GWAS summary statistics^43,44^. Using LDGMs, most of these operations run in ~10 minutes on the entire genome (AFR; 11.6 million SNPs) (Figure 4a; Methods).

**Figure 4:**
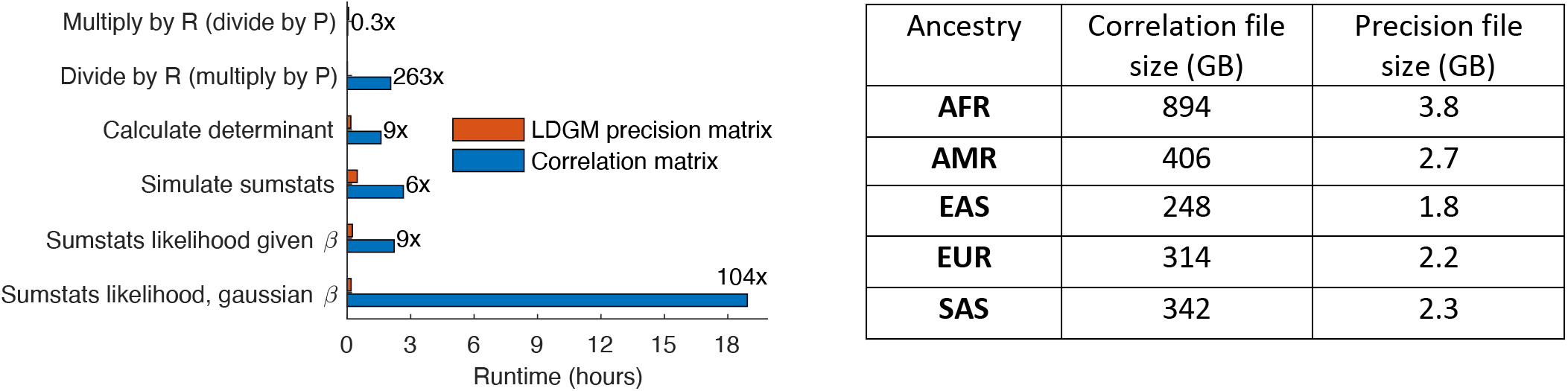
computational advantages of LDGMs. (a) Runtime of matrix operations and likelihood calculations using LDGM precision matrices vs. correlation matrices, for AFR (11.6M SNPs). (b) Total uncompressed file size of LDGM precision matrices vs. correlation matrices in each ancestry group. For numerical results, see Supplementary Table 3.

LDGMs also allow order-of-magnitude reductions in memory usage. The total memory usage of the 1,360 EUR correlation matrices is 314GB (largest: 3.8GB), compared with 2.2GB for the LDGM precision matrices (largest: 6MB) (Figure 4b; Methods). This >100x reduction makes it feasible to work with LD matrices covering the entire genome on a personal computer.

Moreover, memory is often a bottleneck in cluster compute environments, and reduced memory requirements allow for more efficient parallelization.

It is computationally expensive to compute LDGMs and LDGM precision matrices themselves. Computing the LDGM takes an average of ~18 hours per LD block, and computing the LDGM precision matrix takes an average of ~3 hours per block in populations besides AFR (~20 hours in AFR, due to the much larger number of common SNPs) (Supplementary Figure 9). Most users will not have to perform these computations themselves, as we have released precomputed LDGM precision matrices for all five 1000 Genomes populations. Users with access to individual-level GWAS data may compute within-sample LDGM precision matrices by rerunning the precision matrix inference step on precomputed LDGMs (see Discussion).

### Polygenic prediction across populations using LDGMs

A potential application of LDGMs is in methods to calculate polygenic scores (PGS). PGS have gained attention recently for their applications in personalized medicine, but genetic predictors based on pedigrees rather than genotypes have been used in animal breeding for decades, and in fact, sparse precision matrices had an important role in their development. Between 1953 and 1976, Henderson^45–47^ developed the best linear unbiased predictor (BLUP) of phenotypic values (e.g., milk production) from pedigrees, but initially, it seemed necessary to work with a large covariance matrix describing the relationship between every pair of cattle (i.e., the genetic relatedness matrix). This was intractable for large herds, until Henderson showed that the same relationships could be encoded in a sparse precision matrix instead, and the computational bottleneck was lifted.^46,47^ Our use of a sparse precision matrix for LD (i.e., *E*(*X^T^X*)^-1^) parallels Henderson’s for relatedness (i.e., *E*(*XX^T^*)^-1^).

In human datasets with genotypes instead of pedigrees, Henderson’s approach is not directly applicable. However, a SNP-based relatedness covariance matrix can be used instead^48^, and BLUP has also been reformulated to operate on GWAS summary statistics and LD correlation matrices, as implemented in LDpred-inf.^5^ BLUP is not the state of the art for genotype-based prediction, as it imposes an infinitesimal (Gaussian) prior on the distribution of effect sizes, and better performance can be obtained using a non-infinitesimal prior.^5,49^ Nonetheless, BLUP is a simple and canonical genetic prediction method that enables a fair comparison between methods based correlation matrices and methods based on the LDGM. It also provides a starting point for the development of more sophisticated algorithms.

We reformulated BLUP to operate on LDGM precision matrices *(BLUP-ldgm)* and compared it with an implementation that operates on LD correlation matrices *(BLUP-ldcov;* equivalent to LDpred-inf^5^) (see Methods). We applied both methods to UK Biobank summary statistics for four traits (Supplementary Table 6) and 1000 Genomes reference LD; as a gold standard, we also applied *BLUP-ldcov* to within-sample UK Biobank LD (computed by Weissbrod et al.^8^). We calculated the correlation between the resulting PGS weights across chromosomes 21-22 (39 LD blocks). This comparison is much better powered than a comparison of phenotypic prediction accuracy, which would depend on the number of samples in the validation cohort, and it is appropriate for the purpose of comparing *BLUP-ldgm* with *BLUP-ldcov*, which are equivalent except for the LD matrices that are used.

We found that *BLUP-ldgm* was actually closer to the gold standard within-sample weights compared with *BLUP-ldcov* (Figure 5a). Both methods were highly correlated, and the difference was small but highly significant. Since the methods are equivalent, this indicates that the 1000 Genomes LDGM precision matrix is a better approximation to UK Biobank LD than the 1000 Genomes LD correlation matrix. The correlation matrix contains noise (due to the limited number of samples), and by leveraging the independence of haplotypes at different positions, LDGMs usefully regularize the LD matrix. (Concordantly, out-of-sample MSE was higher for the 1000 Genomes correlation matrices than for our 1000 Genomes precision matrices; Supplementary Table 5). We also compared *BLUP-ldgm* and *BLUP-ldcov* with simulated summary statistics and confirmed that their results are nearly identical at small to realistic sample sizes (*N* ≤ 10^6^), becoming divergent at extremely large sample sizes (*N* ≥ 10^7^) (Supplementary Figure 10). These estimates indicate that extremely well-powered GWAS require more accurate LD reference panels (see Discussion), consistent with what has been reported in fine mapping studies.^50^

**Figure 5:**
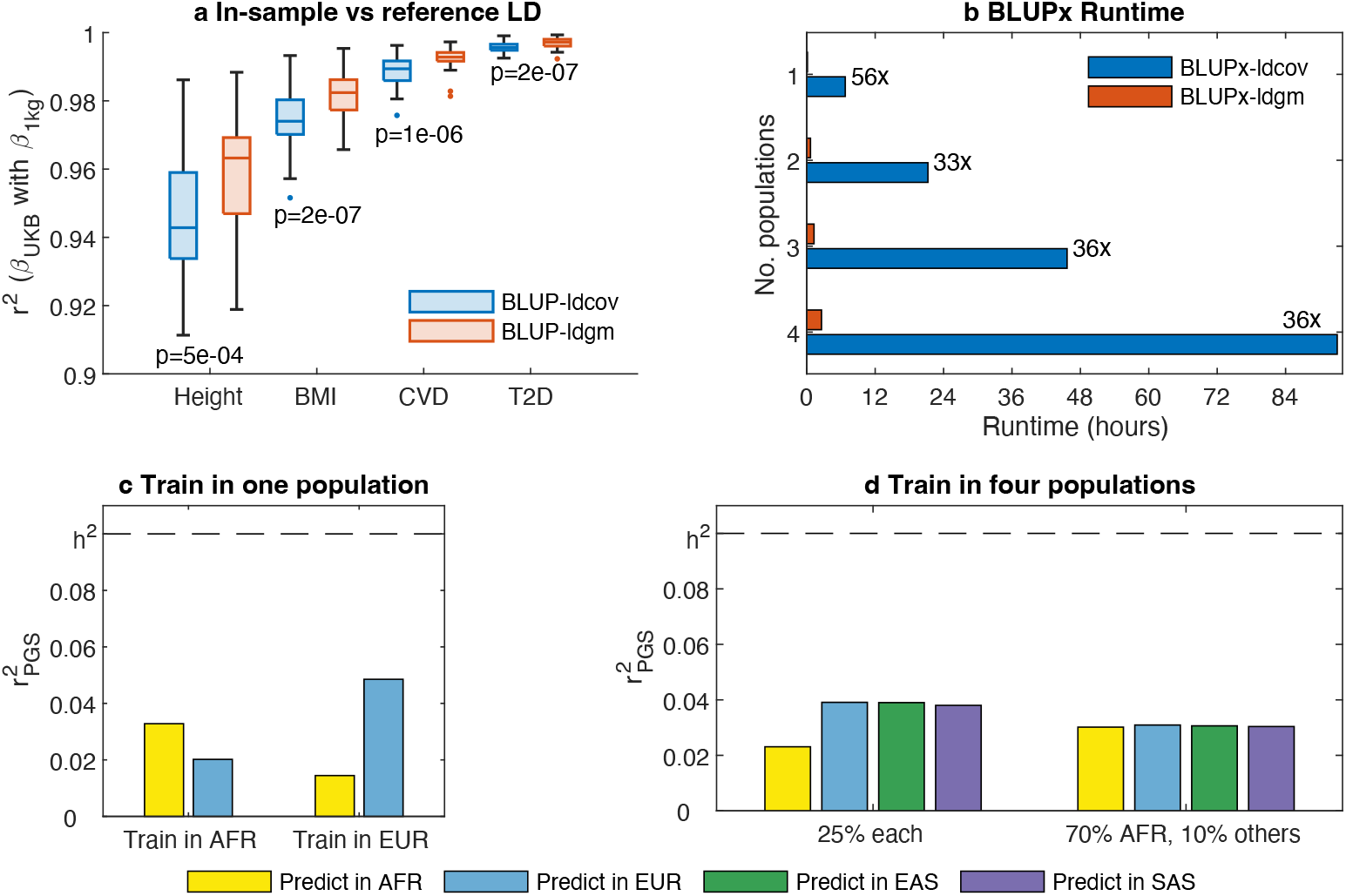
Best linear unbiased prediction with LDGMs. (a) We applied *BLUP-ldgm* and *BLUP-ldcov* to UK Biobank summary statistics for four traits, with 1000 Genomes EUR reference LD, and compared their PGS weights with gold-standard weights calculated from within-UKB LD (using *BLUP-ldcov)*. Box plots indicate the distribution of *r^2^* values across LD blocks (chromosomes 21-22), and binomial p-values are reported. For numerical results, see Supplementary Table 5. (b) Runtime of *BLUPx-ldgm* and *BLUPx-ldcov* in whole-genome simulations with different numbers of ancestry groups. For numerical results, see Supplementary Table 7. (c) Prediction accuracy 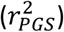 of *BLUP-ldgm* when training in a single ancestry group (EUR or AFR). For numerical results, see Supplementary Table 8. (d) Prediction accuracy of *BLUPx-ldgm* when training in four ancestry groups (AFR, EUR, EAS, SAS), with different relative sample sizes and a total sample size of 1e6. For numerical results, see Supplementary Table 8.

A well-known challenge is that PGS are not portable across genetic ancestries, primarily due to differences in allele frequency and LD, motivating the development of methods that analyze ancestrally diverse GWAS.^6,9,51,52^ BLUP has been extended to handle multiple ancestry groups, but computational complexity is a challenge.^11,53^ We implemented multi-ancestry BLUP using LDGMs (*BLUPx-ldgm*) or covariance matrices (*BLUPx-ldcov*) and compared their runtime in simulations involving one to four populations and the entire genome (1361 LD blocks). *BLUPx-ldgm* was ~40 times faster, running in 2.6 vs. 93 hours in the simulation with four groups (Figure 5b).

Then, we applied *BLUPx-ldgm* to summary statistics simulated from different genetic ancestries in order to quantify non-portability. We simulated summary statistics from their asymptotic sampling distribution^43,44^ with allele frequencies and LDGM precision matrices from AFR or EUR, applied *BLUPx-ldgm* to data from one group or the other, and evaluated its prediction accuracy in each group (see Methods). As expected, scores trained in EUR had much lower accuracy in AFR (concordant with what is observed with real phenotypes^9^). However, non-portability was directionally asymmetric: the decrease in accuracy was smaller when training in AFR and testing in EUR (Figure 5c). Asymmetric non-portability was also observed in other population pairs involving AFR, but it was not observed in non-AFR population pairs (Supplementary Table 8). The apparent asymmetry in PGS portability might be a consequence of the out-of-Africa bottleneck, which reduced the diversity of haplotypes that are found outside of Africa; polygenic prediction in AFR is fundamentally harder than in other ancestry groups, owing to its greater diversity.

Finally, we simulated population specific summary statistics from four ancestry groups (AFR, EUR, EAS, SAS), applied *BLUPx-ldgm* to all of them jointly, and evaluated its prediction accuracy in each. With equal sample size (and heritability) in each ancestry group, prediction accuracy was not equal. Instead, it was ~30% lower in AFR (Figure 5d and Supplementary Table 8), probably also because of greater genetic diversity within AFR. In order to equalize prediction accuracy across groups in these simulations, it was necessary for 70% of the training data to be AFR (10% each EUR, EAS and SAS) (Figure 5d and Supplementary Table 8).

These simulations indicate that LDGMs can be used to greatly improve the computational performance of PGS methods without any loss in accuracy, and that a particularly promising application is the development of PGS methods that are able to handle data from multiple ancestry groups. However, for AFR in particular, large amounts of new data will be required to equalize PGS accuracy, even with the use of multi-ancestry methods.

## Discussion

Computational and statistical challenges in human genetics can be overcome using explicit genealogical models. The Li-Stephens model^15^ is widely used for imputation^54^ and phasing,^55^ and it is also an important component of scalable genealogy inference methods like *tsinfer* and *relate*^18,19^. Large-scale genealogies, in turn, have enabled a variety of powerful methods in statistical and population genetics^26,27,56^. In this study, we leveraged genome-wide genealogies to derive LDGMs and to address computational challenges associated with LD and ancestral diversity.

LDGMs have several promising applications. In polygenic prediction, state-of-the-art Bayesian methods^6,41,49,57,58^ rely on slow LD matrix operations, and scaling across genetic ancestries remains an important challenge despite recent progress^6,41,57^. We have implemented BLUP with LDGMs, achieving large improvements in runtime; BLUP is not the state of the art for polygenic prediction (due to its reliance on an infinitesimal prior), but it is a logical step toward the development of more sophisticated methods. Another promising application is heritability partitioning, where the state-of-the art method, S-LDSC^3,59^, is more widely used than likelihood-based estimators like RSS^4^ due to its speed and scalability, despite likelihood-based methods having better accuracy^39^. A potential application is fine mapping, although we caution that accurate fine mapping may require extremely accurate LD matrices, and individual-level estimation may be the most robust solution.^8,50^

When modeling LD, it is sometimes critical to distinguish between within-sample LD (i.e., the sample covariance matrix of the genotypes), reference LD (i.e., the sample covariance matrix of a different set of genotypes from the same ancestry group), and population LD (i.e., the covariance matrix of the entire population). The LDGM precision matrix is an estimator of population LD; it is based on a model for the conditional dependencies that arise when sampling ancestral haplotypes to produce a new genome, not the particular genomes observed in the study. As such, it should not be used in situations where within-sample LD is strictly needed, even if it is inferred within-sample. Still, LDGM precision matrices inferred within-sample may have advantages over those inferred from a reference panel (especially a relatively small one like 1000 Genomes), as they avoid mismatch between the respective genetic ancestries. When individual level data is available, it may still be advantageous to use LDGMs and summary association statistics because the computational complexity of individual-level methods scales with the size of the study. When individual level data is unavailable, it is possible to assess whether a reference LD matrix is appropriate using DENTIST^60^.

Even though LDGM precision matrices can be used to model ancestry-specific LD patterns, our approach does not rely conceptually on populations as discrete or essential features of human ancestry.^61^ Instead, the LDGM is shared across the ancestries represented in 1000 Genomes. Haplotype frequencies differ with ancestry, but ancestral haplotypes themselves are mostly shared, and it is the haplotypes, not their frequencies, that determine the LDGM. In contrast, LDGM precision matrices do depend on haplotype frequencies, and ancestry groups (especially “EUR”) can be used to match the study population with an appropriate LD reference panel; this heuristic is useful because of the way that most existing association studies are ascertained.

Adjacencies in the LDGM correspond to genealogical, not biological, relationships. Although LD patterns are strongly affected by negative selection, and they differ between functional and neutral regions of the genome, we do not see a reason that LDGMs would be functionally informative for individual variants or clusters of variants. Conversely, statistical methods to analyze GWAS data using LDGMs are not shielded from the effects of LD dependent architecture, which can be an important source of bias if not modeled appropriately. ^59,62^

Our work has several limitations. First, LDGMs do not model population structure or admixture, and they can perform poorly in groups with strong population structure. Our results support their use within the ancestry groups besides AMR defined in 1000 Genomes, but currently, we recommend that they not be applied to admixed populations. Local ancestry inference is a possible approach to address this limitation.^34^ Second, LDGMs and LDGM precision matrices are computationally expensive to compute. This is a barrier for users who wish to tailor LDGM precision matrices to their own genotype data; however, we have released 1000 Genomes LDGM precision matrices for each superpopulation (see Data Accessibility), and we envision that most users will be able to use this resource and avoid the upfront cost. Third, some applications – especially fine mapping – may require more accurate LDGM precision matrices than it is possible to infer from 1000 Genomes. LDGM precision matrices can be tailored to specific datasets, and parameter choices permit a tradeoff between sparsity and within-sample accuracy.

Despite these limitations, LDGMs address statistical and computational challenges associated with genetic association studies and ancestral diversity. We advocate for GWAS consortia and biobanks, especially ancestrally diverse studies without a well-matched reference panel, to publicly release LD information together with their summary association statistics, allowing them to be analyzed using LDGMs.

## Methods

### Dataset preparation and tree sequence inference

We analyzed VCF files containing 1000 Genomes Data sequenced to 30x coverage by the New York Genome Center (see Data Availability).^30^ This phased genetic dataset is in the GRCh38 genome build and contains 3,202 sampled individuals; we analyzed 2,504 approximately unrelated individuals from phase three of the 1000 Genomes project.

We inferred tree sequences from the autosomes of the 2,504 unrelated individuals using tsinfer version 0.2^31^. We first converted the downloaded VCF files to the *.samples* format used by *tsinfer*. We used all biallelic SNPs with high confidence ancestral states in Ensembl release 100^63^ to infer tree sequence topologies using tsinfer’s default parameter settings. This meant that no mismatch parameters were specified, enforcing the infinite-sites assumption. Afterwards, the tree sequences contained 63M SNPs.

However, restricting our analysis to biallelic SNPs with high-confidence ancestral states excluded many SNPs that we wished to incorporate in downstream analyses. Thus, we mapped SNPs with a low-confidence ancestral state or no inferred ancestral state, as well as indels, back onto the tree sequences using parsimony. Our LDGM creation algorithm does not currently support recurrent mutation, so we only kept variants where a single mutation captured the pattern of variation at that site. After adding these variants, our tree sequences contain 93.1% of a comparison set of 7,742,161 imputed, common GWAS SNPs (>1% MAF).

### Generating LDGMs and precision matrices

We estimated precision matrices from 1,361 approximately independent LD blocks and each 1000 Genomes continental ancestry group (see Data Availability).^32^

We used *ldgm* version 0.1 to create LDGMs from the inferred tree sequences with the following parameters: *path_threshold=8, MAF=0.01, recombination_threshold=0.01*. The minor allele frequency threshold was applied such that SNPs with at least a 1% MAF in any of the five ancestry groups were retained in the final LDGM (18M SNPs).

Finally, precision matrices were estimated for each 1000 Genomes ancestry group using the LDGMs from the previous step, using a path threshold of 4 and an L1 penalty of 0.1. In AFR, which has a larger number of common SNPs per block, we used an L1 penalty of 0.2 for some of the blocks in order to reduce runtime (Supplementary Table 1).

### Mean squared error

We define the mean squared error (MSE) between an LD correlation matrix *R* and an LDGM precision matrix *P* as:

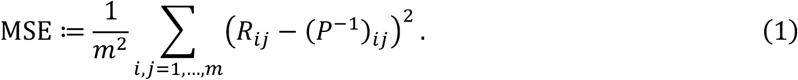

This is proportional to the mean squared error when multiplying a vector by *R* or instead dividing it by *P*, and it is mostly sensitive to the large eigenvalues of *R* (small eigenvalues of *P*). If *x* ~ *N*(0, *m*^-1^*I*), then

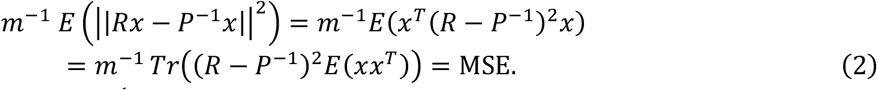

The MSE quantifies how close *P*^-1^ is to *R*. An alternative definition, which quantifies how close *PR* is to the identity matrix, is defined in the Supplementary Note.

### Evaluating LDGMs on simulated and inferred tree sequences

To evaluate the impact of tree sequence inference accuracy on the accuracy and sparsity of LDGM precision matrices, we compared the results of our pipeline on true, simulated tree sequences compared to tree sequences inferred with *tsinfer*. We first used *msprime* to simulate ten replicates of a tree sequence that approximates the size of a typical LD block in our analysis. We simulated 3 Mb of sequence, with 5008 sampled chromosomes, a mutation rate of 1.2e-8 mutations per-base pair per-generation, a recombination rate of 1e-8 recombinations per-base pair per-generation and an effective population size of 10,000. We then used the genotype data from these simulations to infer tree sequences using *tsinfer*.

We ran our LDGM inference pipeline on the simulated and inferred tree sequence from each simulation replicate and evaluated the accuracy of the LDGM precision matrix compared to the correlation matrix (MSE) as well as the sparsity (average degree). The accuracy results are visualized in Figure 3d, and the full results are in Supplementary Table 2.

### Runtime and memory usage

Runtime experiments were performed on the Broad Institute compute cluster, parallelized and summed across LD blocks, using MATLAB R2019a.

Correlation matrix memory usage in bytes was calculated as 8*m*^2^, where *m* is the number of SNPs with AF>0.01 in an LD block. Precision matrix memory usage was calculated as 16*md* + 8(*m* + 1), where *d* is the average degree (and *md* is the number of nonzero entries), which is the minimum memory requirement in MATLAB.

### Best linear unbiased predictor

The BLUP PGS weights have a simple formula:

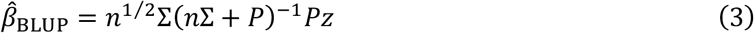

where *n* is the GWAS sample size, Σ is the prior covariance matrix of the per-s.d. effect sizes *β* (e.g., 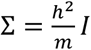), *P* is the LDGM precision matrix, and *z* is the vector of GWAS Z scores. In *BLUP-ldgm*, this formula is evaluated by solving a sparse system of equations:

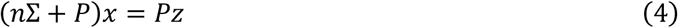

and setting 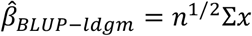. If SNPs in the LDGM are missing from the summary statistics, it is handled as described in the Supplementary Note. *BLUP-ldcov* solves a dense system of equations instead:

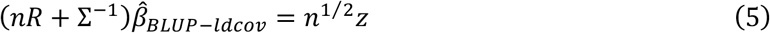

where *R* is the sample covariance matrix. Both *BLUP-ldcov* and *BLUP-ldgm* are parallelized across LD blocks.

BLUPx is a straightforward generalization of BLUP. The Z scores are concatenated across populations, and the precision matrices (or correlation matrices) are concatenated to form a block-diagonal matrix. The prior covariance matrix Σ has nonzero off-diagonal entries corresponding to the same SNP in two different populations. For example, if SNP *j* is present in populations 1 and 2 with effect size variance 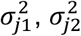 respectively, and the cross-population genetic correlation is *r_pop_*, then the corresponding entry is *σ*_*j*1_*σ*_*j*2_*r_pop_*. If SNP *j* is absent in population 2, then there is no corresponding entry. In our simulations, the effect-size variance for every SNP is the same, and it is chosen to produce the desired heritability in each ancestry group. We produce the cross-population effect-size covariance matrix, concatenate the population-specific precision matrices to form a block-diagonal matrix, concatenate the Z scores to form a longer vector; then, we substitute these matrices into equation (4).

### Polygenic scores with multiple ancestry groups

In our PGS simulations, first, we simulated causal effect sizes from a multivariate normal distribution for every SNP with allele frequency at least 1% in any of the four ancestry groups (AFR, EAS, EUR, SAS). (AMR was excluded due to high its MSE). The effect size variance for each population was chosen so that the expected heritability would be 0.1, and the effect size correlation between each pair of populations was 0.9. For each ancestry group, we simulated summary statistics from their asymptotic sampling distribution based on the LDGM precision matrix and the 1000 Genomes allele frequencies (see Supplementary Note). We applied *BLUPx-ldgm* to these summary statistics, using as a prior the same distribution that was used to simulate the data. Supplementary Table 8 lists the sample sizes that were used in each simulation. To calculate the prediction accuracy, we used the following formula:

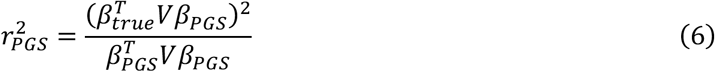

where *β_true_* is the vector of true per-allele effect sizes, *β_PGS_* is the vector of estimated per-allele effect sizes, and *V* is the genotype covariance matrix for the target population (i.e., *DRD* where *D* is a diagonal matrix).

## Acknowledgements

We are grateful to Daniel J. Weiner, Ajay Nadig, Alkes L. Price, Raymond Walters, Hilary Finucane, Xihong Lin, Hui Li, Brieuc Lehmann, Peter Ralph, Gregor Gorjanc, Jerome Kelleher, and Rahul Mazumder for helpful discussions.

## Data accessibility

Tree sequences, LDGMs, and LDGM precision matrices are available at https://github.com/awohns/ldgm_paper. High-coverage phased 1000 Genomes genotype data is available at http://ftp.1000genomes.ebi.ac.uk/vol1/ftp/data_collections/1000G_2504_high_coverage/working/20201028_3202_phased/. LD-independent blocks are available at https://github.com/jmacdon/LDblocks_GRCh38. UK Biobank summary statistics and LD is available at https://alkesgroup.broadinstitute.org/. Pan-UKB LD is available at http://pan.ukbb.broadinstitute.org/.

## Code accessibility

We have released an open-source software package, *ldgm* version 0.1, implemented in python and MATLAB. *ldgm* allows inference of LDGMs and LDGM precision matrices, as well as computationally efficient analyses of GWAS summary statistics using LDGMs. It is available at https://github.com/awohns/ldgm. All of the functions for analyzing GWAS summary statistics with LDGMs, including *BLUPx-ldgm*, are currently implemented in MATLAB; a Python implementation is planned. Scripts to reproduce the results of this manuscript are available at https://github.com/awohns/ldgm_paper.

## Author contributions

A.W.W., P.S.N. and L.J.O. developed the methods. E.S.L., B.M.N. and A.B. suggested analyses. A.W.W., P.S.N., J.L.B. and L.J.O. performed the experiments. A.W.W., P.S.N., J.L.B., B.M.N., and L.J.O. wrote the paper. L.J.O. supervised the research.

