## Supplementary figures and table captions for "Extremely sparse models of linkage disequilibrium in ancestrally diverse association studies"

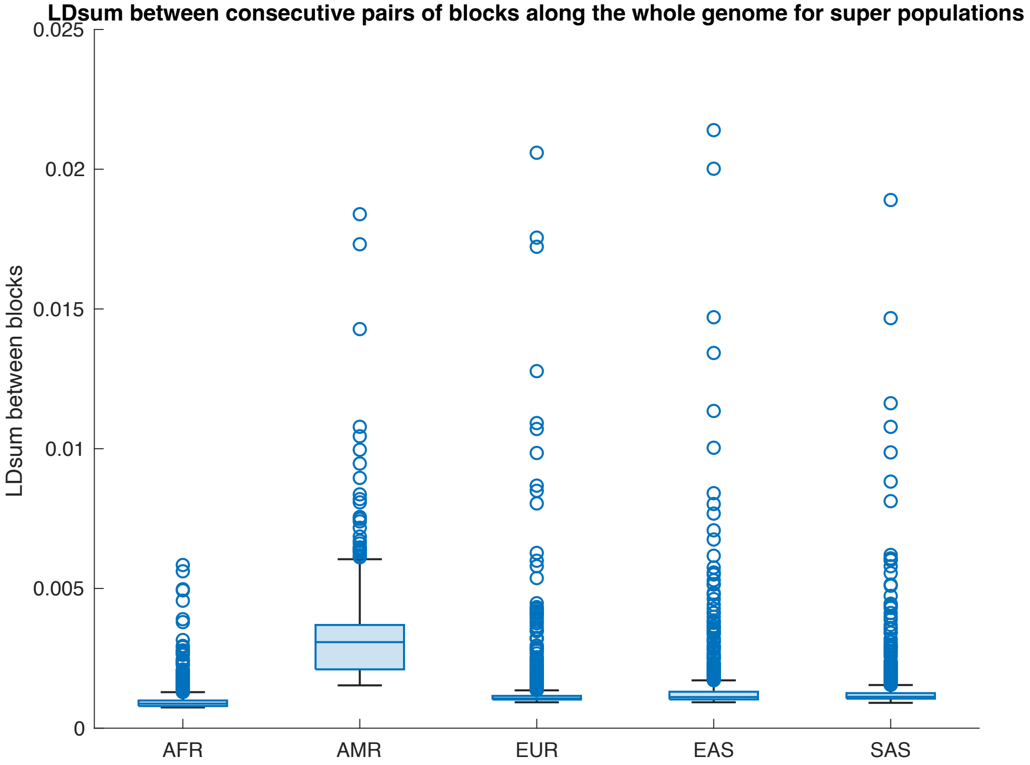


**Supplementary Figure 1: LD between adjacent pairs of LD blocks.** For all blocks of chromosomes 1 to 22, we evaluated the mean $r^2$ between every pair of SNPs ("LDsum") in consecutive LD blocks, within each ancestry group. The expected mean $r^2$ is around 1e-3, i.e. $1/2n$.

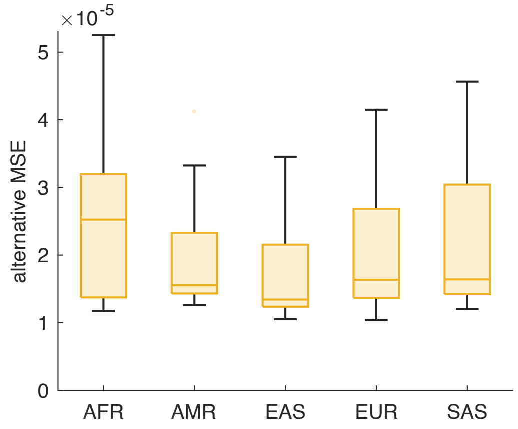


**Supplementary Figure 2: accuracy of LDGM precision matrices by an alternative error metric.** The alternative MSE measures the difference between $PR$, the product of the LD correlation matrix and the LDGM precision matrix, and the identity matrix (see Supplementary Note). Compared with the MSE, the alternative MSE is less sensitive to large eigenvalues of $R$, probably explaining why it is not elevated for AMR. We show the distribution of within-sample altMSE values across LD blocks on chromosome 22, for each 1000 Genomes ancestry group.

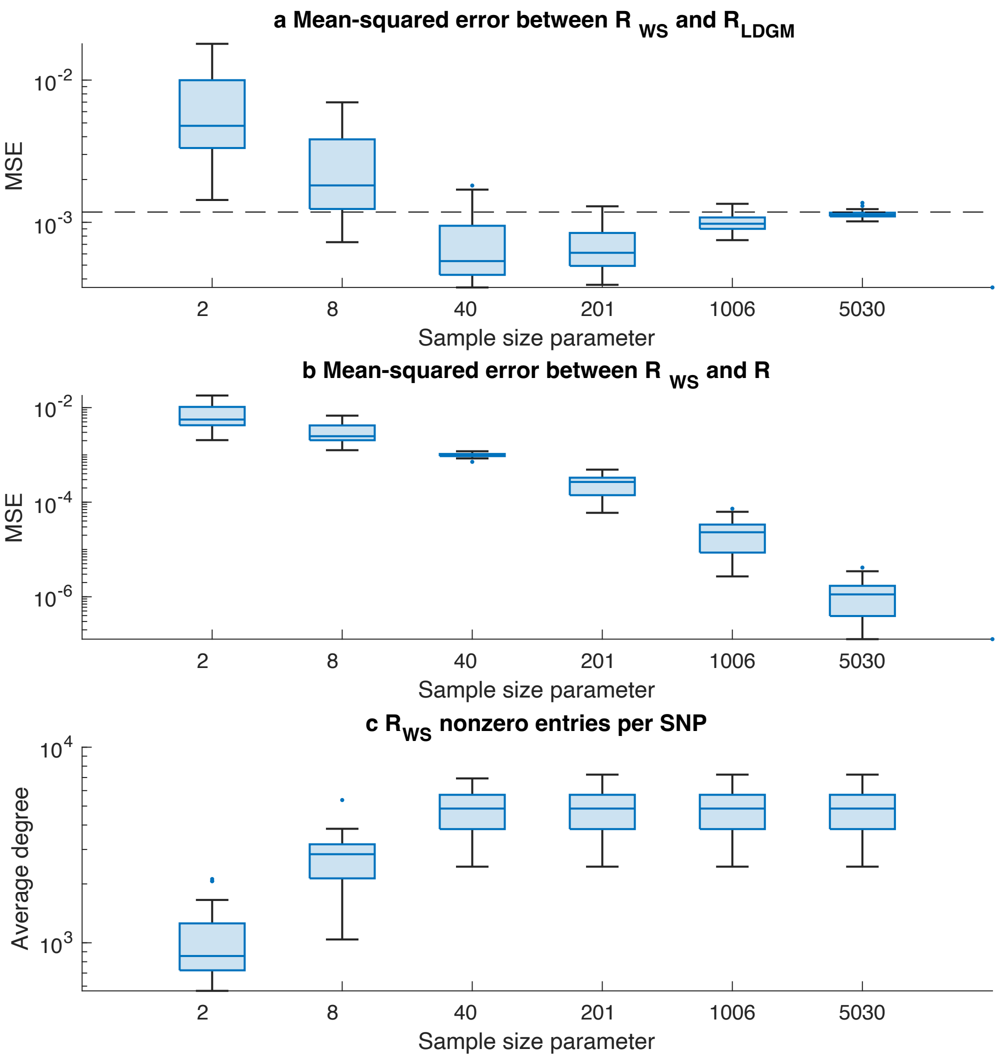


**Supplementary Figure 3: comparison of LDGM precision matrices with Wen-Stephens shrinkage estimator.** The comparison was performed in EUR, on chromosome 22 only (20 LD blocks). To vary the amount of shrinkage, we changed the sample size parameter in the Wen-Stephens estimator (actual sample size: 1006). (a) Mean-squared error between the Wen-Stephens estimator and the LDGM precision matrix inverse. Dotted line denotes the median MSE between the LD sample correlation matrix and the LDGM precision matrix inverse. (b) Mean-squared error between the Wen-Stephens estimator and the sample correlation matrix. Values are larger than the corresponding numbers in panel (a) for sample size parameters up to 40, and smaller for sample size parameters of 201 or higher. (c) Number of nonzero entries per SNP in the Wen-Stephens estimator. Correlations with absolute value less than 1e-8 are set to zero (consistent with the original paper), resulting in slightly increased sparsity for small values of the sample size parameter. At larger parameter values, no SNP pairs are below the threshold within LD blocks, but this approach can still be used to produce a sparse, banded diagonal matrix when it is not desired to use discrete blocks. Somewhat more sparsity can be achieved by relaxing the 1e-8 threshold, but not without causing increased error.

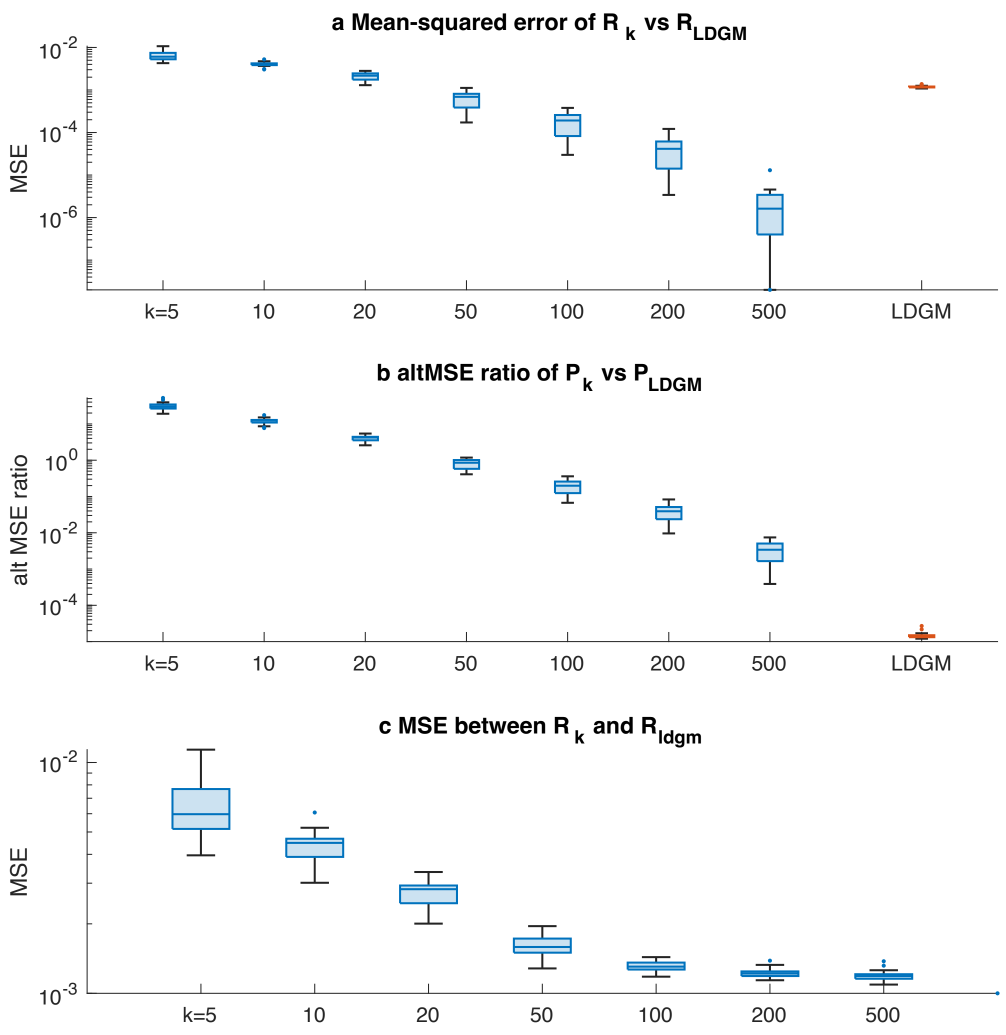


**Supplementary Figure 4: comparison of LDGM precision matrices vs. rank-*k* approximations for the local LD matrix.** The comparison was performed in EUR, on chromosome 22 only (20 LD blocks), and we considered different values of *k*. (a) To quantify the accuracy of the rank-k approximation to the LD correlation matrix, we computed its MSE at different values of *k*. $k = 10$ corresponds most closely to the density of the LDGM precision matrix, which is a symmetric matrix with 20 nonzero entries per SNP (10 per SNP in its upper triangle). The MSE at $k = 10$ was about 3 times higher than that of the LDGM precision matrix; it was most similar at $k = 50$. MSE is always zero when $k$ is greater than or equal to the sample size (i.e., when *k*=1006). (b) To quantify the accuracy of the rank-k approximation to the LD precision matrix, we computed the *alternative MSE ratio*, which quantifies whether the approximate precision matrix multiplied by the correlation matrix is close to the identity (see Supplementary Figure 2 and Supplementary Note). By this metric, the LDGM performs much better than a rank-k approximation even at $k = 500$. (c) We calculated the MSE between the rank-k approximation and the inverse of the LDGM precision matrix. This was never significantly smaller than the MSE of the LDGM precision matrix inverse with the sample correlation matrix.

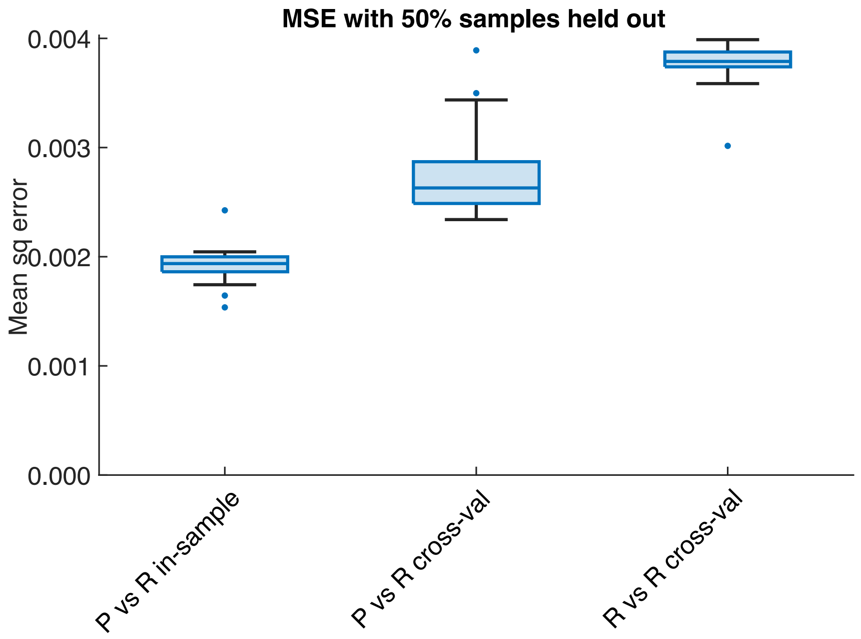


**Supplementary Figure 5: performance of LDGM precision matrices in cross validation.** For each LD block on chromosome 22, we randomly split the 1000 Genomes EUR haploid samples into two subsets of equal size. We computed an LDGM precision matrix from one of the two subsets (the LDGM was constructed from all samples in 1000 Genomes). We computed the MSE for three comparisons: the precision matrix vs. the correlation matrix from the same sample; the precision matrix vs. the correlation matrix from the opposite sample; and the correlation matrix from one sample vs. the correlation matrix from the opposite sample.

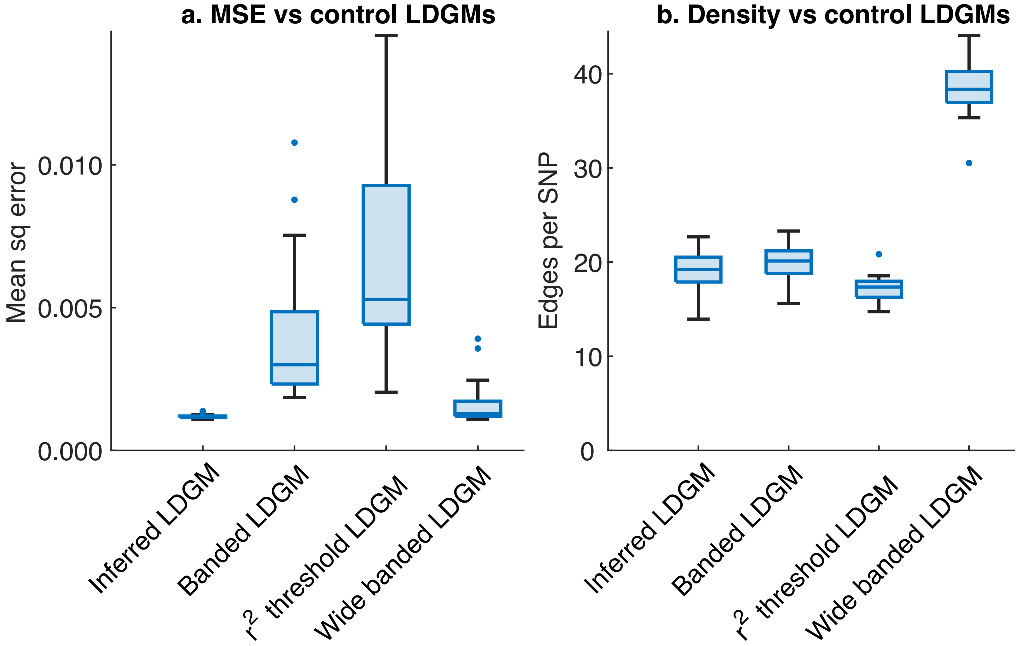


**Supplementary Figure 6: accuracy and sparsity of precision matrices constructed from naïve LDGMs.** In 1000 Genomes EUR data from chromosome 22, we compare our inferred LDGM (derived from tree sequences) with a banded-diagonal LDGM, an $r^2$ threshold LDGM, and a banded-diagonal LDGM with a large band. For the first banded LDGM, the band size was chosen to match the number of edges with path weight less than 4 in our tree-sequence based LDGM for each LD block (approximately 50 edges per SNP). For the r2-threshold LDGM, the threshold was chosen to produce the same number of edges. For the large banded-diagonal LDGM, we used a band size that corresponded to the number of edges with path weight less than 8 (approximately 300 edges per SNP). For each LDGM, we computed precision matrices with an L1 penalty of 0.1, and we calculated the mean squared error (a) and the number of edges per SNP in the precision matrix (b).

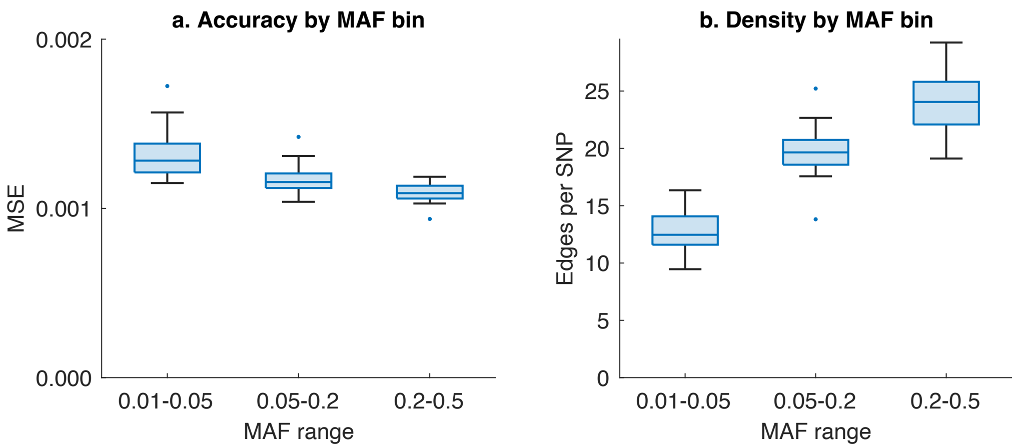


**Supplementary Figure 7: accuracy and density of LDGM precision matrices at different allele frequencies**. For the 20 EUR LDGM precision matrices on chromosome 22, we partitioned SNPs into three bins by their minor allele frequency in EUR. Each bin contained a similar number of SNPs. (a) MSE across pairs of SNPs (i,j) where SNP i has the specified allele frequency (and SNP j may or may not). (b) Average number of neighbors per SNP in each MAF bin (including edges with SNPs not in the bin).

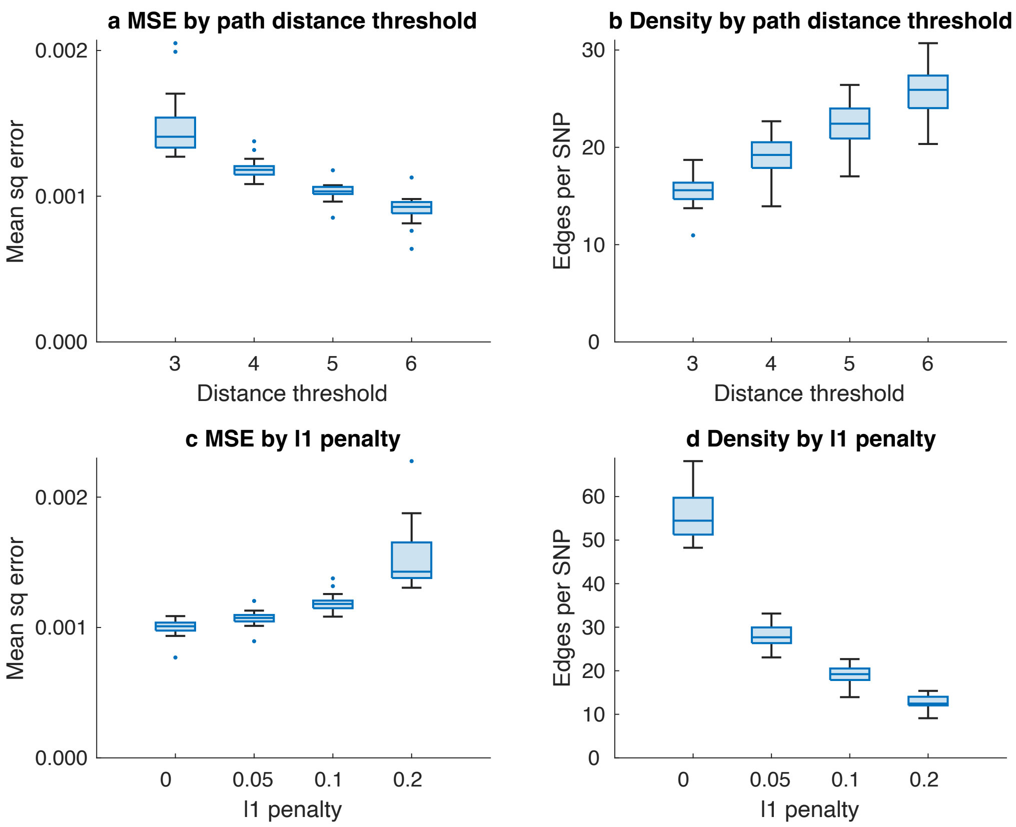


**Supplementary Figure 8: tradeoff between accuracy and sparsity with different parameter settings**. On chromosome 22 EUR data, we varied the path distance threshold (panels a-b) and the L1 penalty (c-d). Our default parameter settings are a distance threshold of 4 and an L1 penalty of 0.1. Precision matrix inference runtime also varies with parameter settings, with greater runtime for settings that produce higher density.

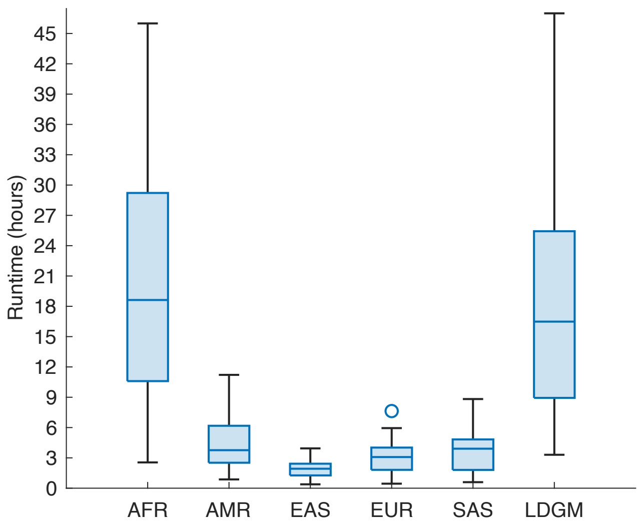


**Supplementary Figure 9: runtime for LDGM and precision matrix inference across chromosome 22.** The first five boxplots indicate the runtime for precision matrix inference for each ancestry group; the sixth indicates the runtime to derive the LDGM from the original tree sequence. For the LDGM inference step, we used 5 compute threads (1 for the precision matrix inference step). Runtime varies across LD blocks and ancestry groups due to variation in the number of SNPs. For numerical results, see Supplementary Table 4.

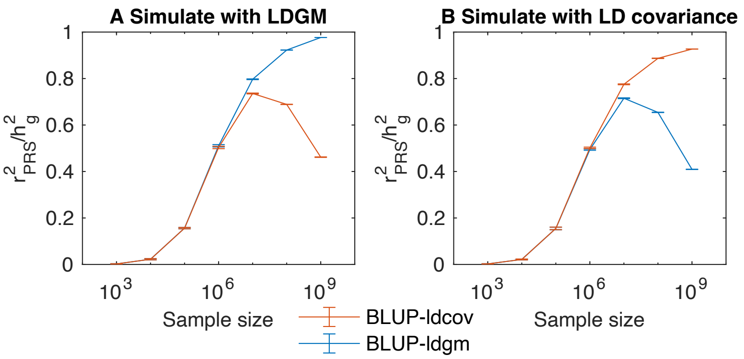


**Supplementary Figure 10: comparison of *BLUPx-ldgm* and *BLUPx-ldcov* in simulations.** We simulated summary association statistics from their asymptotic sampling distribution using either the LDGM precision matrix (a) or the LD covariance matrix (b), at different sample sizes, for the whole genome. *BLUP-ldgm* and *BLUP-ldcov* had similar performance (PRS $r^2$) at small to realistic sample sizes, but at very large sample sizes, better performance was obtained using the LD matrix that matched the way the data was simulated.

**Supplementary Table Captions**

**Supplementary table 1 (see Excel file): statistics for each LDGM precision matrix.** For each ancestry group-LD block pair, we report the number of SNPs, the number of edges per SNP (degree) before and after L1-penalized precision matrix inference, the MSE, the alternative MSE, and the file size. We also list the parameters that were used for inference (the path distance threshold, the L1 penalty, and the recombination frequency threshold).

**Supplementary table 2 (see Excel file): numerical results for Figure 3d.**

**Supplementary table 3 (see Excel file): numerical results for Figure 4a.**

**Supplementary table 4 (see Excel file): runtime per LD block to derive the LDGM and infer the LDGM precision matrix.** Results are reported for chromosome 22 only because for most of the genome, we used an old version of the software that ran more slowly. For the LDGM inference step, we used 5 compute threads (1 for the precision matrix inference step). (a) Runtime for precision matrix inference for each LD block and each population; (b) runtime for LDGM inference for each LD block.

**Supplementary table 5 (see Excel file): numerical results for Figure 5a.** We also report the MSE between the UK Biobank LD correlation matrix and either the 1000 Genomes LDGM precision matrix or the 1000 Genomes LD correlation matrix. (a) Results for each LD block (chromosomes 21-22). We report the MSE between the UK Biobank correlation matrix and either the 1kg correlation matrix or the 1kg LDGM precision matrix. Similarly, we also report the $r^2$ between the BLUP PGS weights, using either the 1kg correlation matrix or the 1kg LDGM precision matrix, for each trait. (b) Averages and comparisons across LD blocks. For the MSE and the BLUP PGS weights $r^2$, p-values are reported for whether the 1kg precision matrix median is significantly better than the 1kg correlation matrix median, calculated using a single-tailed binomial test.

**Supplementary Table 6 (see Excel file): Phenotypes summary association statistics analyzed**. Summary statistics were calculated by Loh et al. (see Data Availability), who also computed heritability and effective sample size.

**Supplementary table 7 (see Excel file): numerical results for Figure 5b.**

**Supplementary table 8 (see Excel file): numerical results for Figure 5c-d.** We report the sample size in each population, the target population, and the prediction $r^2$. We also report cross-ancestry prediction accuracy at smaller sample sizes and for other population pairs.
