## Supplementary note for "Extremely sparse models of linkage disequilibrium in ancestrally diverse association studies"

### 1. Probabilistic Graphical Model of Genotypes

In this section, we precisely define the conditional dependency and independency of SNPs. Then we define two classes of graphs that can represent conditional dependencies and independencies. We will produce a graphical model that represents all conditional dependencies that is minimal under some theoretical assumptions.

Let $S = \{1, 2, \ldots, m\}$ denote a set of SNPs and let $H \subseteq 2^S$ denote a set of haplotypes. We analyze the conditional dependencies among SNPs that arise when sampling one of these haplotypes at random. Initially, we assume that haplotypes in $H$ are observed completely, without missing SNPs; in practice, ancestral haplotypes are only partially observed, and we consider this case at the end of this section. For a subset of SNPs $A$ and a haplotype $h \in H$ the restriction of $h$ to $A$ is denoted by $A(h)$.

Let $\Pi$ be the set of probability distributions with support $H$, let $P \in \Pi$, and let $X$ be a random variable with distribution $P$. A subset of SNPs $A$ is identified with the random variable $X_A = A(X)$ chosen under $P$, to $A$.

For three sets of SNPs $A, B, C$, $X_A$ is conditionally independent of $X_B$ given $X_C$ if their joint probability density factors:

$\forall c \; s.t. P(X_C = c) > 0, \; P(X_A = a, X_B = b \,| X_c = c) = P(X_A = a | X_C = c) \, P(X_B = b | X_C = c).$

Depending on $H$, certain conditional independencies are possible, and others are not:

**Proposition 1.** For three sets of SNPs $A, B, C$, $X_A$ and $X_B$ are conditionally dependent given $X_C$ for some distribution $P \in \Pi$ if and only if for some pair of haplotypes $h_1$and $h_2$, the symmetric difference $h_1 \Delta\, h_2$ overlaps $A$ and $B$ but not $C$.

**Proof:** First, suppose there is no such pair of haplotypes; for any $h_1, h_2$, without loss of generality,

$$P(A = A(h_1)|\, C = C(h_1)) \;\; P(B = B(h_1)|\, C = C(h_1)) = P(B = B(h_1)|\, C = C(h_1))$$
$$= P(A = A(h_1), B = B(h_1)|\, C = C(h_1))$$

Therefore $A$ and $B$ are conditionally independent given $C$.

Second, suppose that for some $h_1, h_2 \in H$, $h_1 \,\Delta\, h_2$ overlaps $A$ and $B$ but not $C$. Let $c = C(h_1)$, and let $P(h_1) = \frac{1}{2}$ and $P(h_2) = \frac{1}{2}$. Then:

$$P(X_A = A(h_1), X_B = B(h_1)|X_c = c) = \frac{1}{2} \neq \frac{1}{4}$$
$$= P(X_A = A(h_1)|X_C = c) \, P(X_B = B(h_1)|X_C = c)$$

■

A simple graph $G = (S, E)$ is an *independency map* (or *I-map*) of $H \subset \{0,1\}^{|S|}$ if for all disjoint subsets $A, B, C$ of $S$ we have

$$A \perp_{\forall P \in \Pi} B|C \;\Leftarrow\; A \perp_G B|C$$

Similarly, $G$ is a *dependency map (or D-map)* of $\Pi$ if

$$A \perp_{\forall P \in \Pi} B|C \;\Rightarrow A \perp_G B|C$$

$G$ is said to be a *perfect map* of $\Pi$ if it is both a D-map and an I-map. These three types of graph were defined for a particular probability distribution in ref [1], and here the definitions are modified to include all dependencies that arise in any $P \in \Pi$.

**Proposition 2:** $G$ is an I-map for $S$ and $H$ if and only if for all $h_1, h_2 \in H$, $h_1 \Delta\, h_2$ is a connected set in $G$.

**Proof:** Suppose that for all $h_1, h_2 \in H$, $h_1 \Delta\, h_2$ is a connected set in $G$. If $X_A, X_B$ are conditionally dependent given $X_C$ for some $P \in \Pi$, then by Proposition 1, there exist $h_1, h_2$ such that $h_1 \Delta\, h_2$ contains some $a \in A, b \in B$ but not any $c \in C$. $h_1 \Delta\, h_2$ is connected, so there is a path in $G$ from $a \in A$ to $b \in B$ contained within $h_1 \Delta\, h_2$; this path does not pass through $C$, so $G$ is an I-map.

Suppose not: for some $h_1, h_2 \in H$, $h_1 \Delta\, h_2$ is not a connected set in $G$. For some $a, b \in h_1 \Delta\, h_2$, every path from $a$ to $b$ in $G$ passes through a set $C$ with $h_1 \cap C = h_2 \cap C$. By Proposition 1, there exists $P \in \Pi$ such that $a, b$ are conditionally dependent given $C$, and $G$ is not an I-map.

■

Not all sets of haplotypes have a perfect map. For example, suppose $S = \{a, b, c\}$ and $H = \{ABC,\ aBC,\ abc\}$. Suppose $G$ is an I-map. By Proposition 2, $bc \in E(G)$ and $\{a, b, c\}$ must be a connected set in $G$. Since $G$ is an I-map, either $ab$ or $ac \in E(G)$ and hence $G$ is not a D-map.

**Problem 1:** Which sets of haplotypes H have a perfect map?

In special cases, a perfect map exists. A set of haplotypes $H$ is called *recombination-free* if there exists a rooted tree $T = (H, E)$ such that the root has no derived alleles and an injection $M: S \rightarrow E$ such that for all SNPs $m \in S$, the haplotypes with the derived allele of SNP $m$ are exactly those that come after $M(m)$ with respect to $T$. Some edges of $T$ have labels associated with mutations i.e., elements of $M(H)$. Define the graph $G(S, E)$ such that two SNPs are adjacent if and only if the unique path connecting them contains no labeled edge. In other words, $G$ is the reduced graph of the line graph of $T$.

**Theorem 1:** Suppose $H$ is recombination-free. Then $G$, the reduced line graph on vertices corresponding to labeled edges of $T$, is the perfect map.

**Proof:** First we show that $G$ is an I-map. Suppose $h$ is the most recent common ancestor of $h_1, h_2 \in H$ (possibly, $h = h_1$). In $T$ there are unique paths $P_1$ and $P_2$ from $h$ to $h_1$ and $h_2$ respectively. Let $S_i$ be the set of labeled edges of $P_i$ for $i = 1, 2$. Then $h_1 \Delta\, h_2 = S_1 \cup S_2$, because $h_1$ has the derived allele for SNPs $S_1$ and $h_2$ has the derived allele for SNPs $S_2$. By the construction of $G$, $S_1 \cup S_2$ is connected; therefore, by Proposition 2, $G$ is a I-map.

Now we show that $G$ is a D-map. Suppose $a, b \in S$ and $C \subset S$. Suppose the unique path in $T$ between $a$ and $b$ does not intersect $C$ i.e., $a$ and $b$ belong to the same connected component of $G \setminus C$. We will show that $a$ and $b$ are conditionally dependent given $C$. There are two cases, $a$ and $b$ are either on the same branch with respect to the root of $T$ or not.

**Case 1.** Suppose $a$ and $b$ are on the same branch of $T$. Let $h_1$ be the haplotype before $a$ and $h_2$ be the haplotype after $b$. Then $h_1 \Delta h_2$ has both $a$ and $b$ but not any $c \in C$ since any $c \in C$ either belong to both or neither of $h_1$ and $h_2$ depending on whether $h_1$ has $c$ or not, and by Proposition 1, $a$ and $b$ are conditionally dependent given $C$.

**Case 2.** Suppose $a$ and $b$ are on different branches of $T$. Let $h$ be the MRCA haplotype of $a$ and $b$. Let $h_1$ and $h_2$ be direct descendant haplotypes of $a$ and $b$, respectively. Then $h_1 \Delta h_2$ has both $a$ and $b$ but not any $c \in C$ since any $c \in C$ either belong to both or neither of $h_1$ and $h_2$ depending on whether $h$ has $c$ or not, and by Proposition 1, $a$ and $b$ are conditionally dependent given $C$.

■

**Problem 2:** For a given set of Haplotypes $H$ with SNPs set $S$ find a sparse simple graph $G = (S, E)$ that is an I-map.

Using Proposition 2, a sufficient condition for $G$ is that for all pairs of haplotypes $h_1, h_2 \in H$, $h_1 \Delta\, h_2$ forms a connected set. In the next section, we define a simple graph called the LDGM and prove that the symmetric difference of any two haplotypes is a connected set of LDGM. Additionally, the LDGM is edge-minimal under a precise condition.

**Missing SNPs.** In practice, ancestral haplotypes are only partially observed. We generalize the definition of conditional dependency and prove Proposition 1 in the generalized setting.

Let the set of haplotypes be $H \subseteq \{0,1,-1\}^m$ where for $h \in H$, $h_i = -1$ if $h$ is missing at position $i$. First, we generalize the symmetric difference to this setting. For $h_1, h_2 \in H$ let their symmetric difference comprise the SNPs where they are different and non-missing:

$$h_1 \,\Delta\, h_2 = \{s \in S\text{: either } s(h_1) = 1, s(h_2) = 0 \text{ or } s(h_2) = 0, s(h_1) = 1\}.$$

We write that $h_1 \equiv h_2$ when $h_1 \,\Delta\, h_2 = \emptyset$, i.e. when $h_1, h_2$ are identical at every position where they are both non-missing.

Next, let $\Pi$ be a set of probability distributions with support $H$, let $P \in \Pi$, and for a subset of SNPs $A$, let $X_A$ be a random variable with distribution $P$ restricted to SNPs in $A$. Recall that for a haplotype $h \in H$, by $A(h)$ we mean haplotype $h$ restricted to $A$. For a second set of SNPs $B$, let

$$H_B = \{h \in H: -1 \notin B(h)\}$$

and let

$$H_B(A, a) = \{h \in H_B: A(h) \equiv a\}.$$

We define:

$$P_B(X_A \equiv a) = \frac{\sum_{h \in H_B(A,a)} P(h)}{\sum_{h \in H_B} P(h)}$$

and likewise for e.g., $P_B(X_A \equiv a \mid X_C \equiv c)$.

This notation allows us to define conditional dependency between two sets of SNPs in the presence of missingness, by effectively conditioning on non-missingness. For three disjoint sets of SNPs $A, B, C$, we say that $A$ and $B$ are conditionally dependent given $C$ if for some $c = C(h)$, there exist subsets $A' \subset A$ and $B' \subset B$ with $a = A'(h)$ and $b = B'(h)$ such that

$$P_{A' \cup B'}(X_{A'} \equiv a, X_{B'} \equiv b \mid X_C \equiv c) \neq P_{A' \cup B'}(X_{A'} \equiv a | X_C \equiv c)\, P_{A' \cup B'}(X_{B'} \equiv b | X_C \equiv c).$$

The reason $A'$ and $B'$ are needed, instead of $A$ and $B$, is explained by this example. Let $h_1(A) = (0)$, $h_1(B) = (0{,}0)$, $h_2(A) = (1)$, and $h_2(B) = (1, -1)$. With our definition, $A$ and $B$ can be conditionally dependent with $B'$ comprising the first element of $B$; however, $h_2 \notin H_{A \cup B}$, so $h_2$ would not produce a conditional dependency if dependencies were defined in terms of $P_{A \cup B}$.

We generalize Proposition 1:

**Proposition 3 (generalizing Proposition 1):** For three sets of SNPs $A, B, C$, $X_A$ and $X_B$ are conditionally dependent given $X_C$ for some distribution $P \in \Pi$ if and only if for some pair of haplotypes $h_1$ and $h_2$, the symmetric difference $h_1 \,\Delta\, h_2$ overlaps $A$ and $B$ but not $C$.

**Proof:** First, suppose that for all possible values $c$ of $C(h)$, for all $h_1, h_2 \in H_\emptyset(C, c)$, $h_1 \,\Delta\, h_2$ is disjoint from either $A$ or $B$. For any such $c$, without loss of generality suppose $h_1 \,\Delta\, h_2$ is disjoint from $A$; let $A' \subset A$ and $B' \subset B$, and let $a'$ be $a$ restricted to $A'$. Then

$$P_{A' \cup B'}(X_{A'} \equiv a' | \, X_C \equiv c) = 1$$

because $A(h) \equiv a$ implies $A'(h) \equiv a'$. It follows that

$$P_{A' \cup B'}(X_{A'} \equiv a | X_C \equiv c)\, P_{A' \cup B'}(X_{B'} \equiv b | X_C \equiv c) = P_{A' \cup B'}(X_{B'} \equiv b | X_C \equiv c)$$
$$= P_{A' \cup B'}(X_{A'} \equiv a, X_{B'} \equiv b | X_C \equiv c)$$

and $A$ and $B$ are conditionally independent given $C$.

Second, suppose that for some $h_1, h_2 \in H$, $h_1 \,\Delta\, h_2$ overlaps $A$ and $B$ but not $C$. Let $c = C(h_1)$, let $A' = A \cap (h_1 \Delta\, h_1) \neq \emptyset$ and $B' = B \cap (h_1 \Delta\, h_1) \neq \emptyset$. Let $P(h_1) = \frac{1}{2}$ and $P(h_2) = \frac{1}{2}$. Then:

$$P_{A' \cup B'}(X_{A'} \equiv A'(h_1), X_{B'} \equiv B'(h_1) | X_C \equiv c) = \frac{1}{2} \neq \frac{1}{4}$$
$$= P_{A' \cup B'}(X_{A'} \equiv A'(h_1) | X_C \equiv c)\, P_{A' \cup B'}(X_{B'} \equiv B'(h_1) | X_C \equiv c).$$

■

### 2. LD Graphical Models

In this section, we generalize the approach in Theorem 1 to the case of recombination. We define a tree sequence, modify the tree sequence so that the graph reduction of Theorem 1 can be applied, and we show that the resulting graph is an I-map (Theorem 2.1) satisfying a minimality condition (Theorem 2.2). We illustrate each step of the algorithm with an example (Example 1).

#### 2.1. Tree Sequences

We provide a formal definition of a *tree sequence*[2]:

**Definition 2.1.** A tree sequence is a directed acyclic multigraph $T$ whose vertices are a set of ancestral and sample haplotypes H, with the following properties:

- Each edge of $T$ is a tuple $(h_1, h_2, S)$, where $h_1$is the "parent haplotype", $h_2$ is the "child haplotype", and $S$ is the set of positions that are spanned by that edge
- For every position $n$, the set of haplotypes that exist at that position and the set of edges $(h_1, h_2, S)$ with $n \in S$ form a rooted tree.
- Every mutation is identified with a position $n$ and a clade of the rooted tree at position $n$; the haplotypes that have that mutation are precisely those contained in that clade, and haplotypes that do not have mutation are those that exist at that position but are not in the clade.

SNPs are encoded in the tree sequence as mutations on specific edges. Ordinarily, the carriers of a SNP may depend not only on which edge this is but also on its position within that edge.

**Example 1.A. (Example from Figure 1)** The tree sequence below has 5 sample haplotypes and 5 ancestral haplotypes. The tree sequence has two positions, each with two mutations. Haplotypes 5 and 6 recombined in haplotype 2.

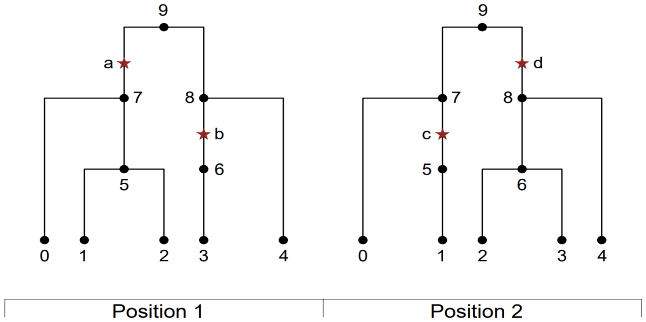


**Example 2. (Tree sequence with no perfect map)** The tree sequence below has the same genealogy as Example 1.A, but different mutations. Suppose for contradiction that $G$ is a perfect map. Haplotype 5 has mutations $\{a, b\}$ and haplotype 2 has mutations $\{a, b, c, d, e\}$. Therefore their symmetric difference is $\{c, d, e\}$ that means $\{c, d, e\}$ must form a connected set. The symmetric difference of haplotypes 6 and 9 implies that $\{c, d\}$ is connected, i.e., $cd \in E(G)$. Since the only haplotypes whose symmetric difference has both $c$ and $e$ are 9 and 2, $c$ and $e$ are independent given $d$ which implies that $ce \notin E(G)$. Therefore $de$ must be an edge and hence $d$ is a cut vertex splitting $c$ and $e$. On the other hand, any symmetric difference containing $d$ abd $e$ also intersects $\{a, b, \ c\}$ but $\{a, b, c\}$ is not a cut set splitting $d$ and $e$ which is a contradiction.

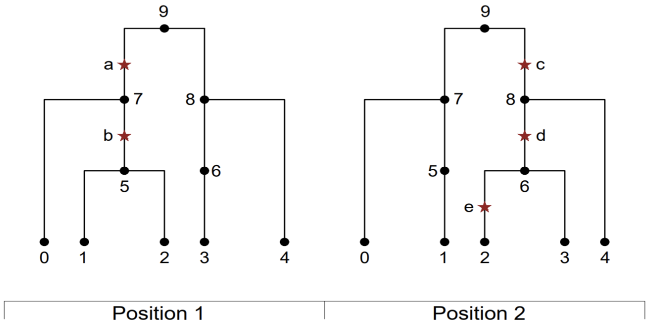


#### 2.2. Bricked tree sequences

In an ordinary tree sequence, if a haplotype has different descendants at each position due to recombination, then a mutation occurring on a single edge would have different carriers depending on its position. This property motivates us to modify the tree sequence, bifurcating edges when they have different descendants at different positions (see section 3.1). We refer to the resulting edges as *bricks*; bricks have the same descendant haplotypes at each position. The resulting graph, which can have multiple edges between the same two haplotypes, is referred to as a *bricked tree sequence*.

**Definition 2.2** A *bricked tree sequence* is a tree sequence with the property that for every edge ("brick") $(h_1, h_2, X)$, the descendent haplotypes of $h_2$ are the same at every position $x \in X$.

**Example 1.B. (Example from Figure 1a)** This example shows the bricked tree sequence. Note that each brick must have a unique set of descendent haplotypes. For example, the edge between haplotypes 8 and 9 has $\{0, 1, 2\}$ and $\{0, 1\}$ descendent haplotype sets in the left and right tree respectively. Therefore, this edge must create two bricks. In the following brick diagram, the corresponding two bricks are represented by assigning two colors (green and purple).

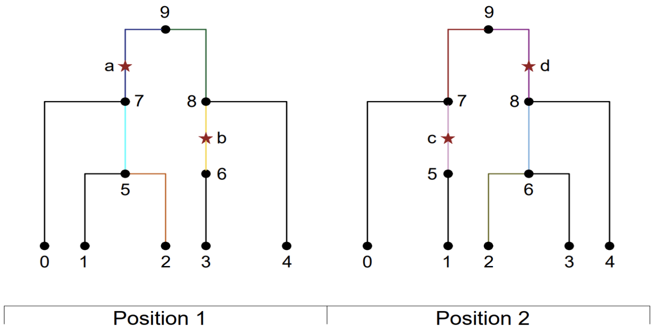


### 2.3 Brick diagrams and the LDGM

In this section, we introduce *brick diagrams,* which describe the symmetric difference between two haplotypes in a tree sequence. We use brick diagrams to define the LDGM, and we show that the LDGM is correct and minimal.

#### 2.3.1 Brick diagrams

In the no-recombination case, the symmetric difference between two haplotypes was the SNPs along the unique path between them in the single tree. A brick diagram is a generalization of a path between two haplotypes.

**Definition 2.3.** The brick diagram of two haplotypes is a subset of the edges of the bricked tree sequence. A brick is in the brick diagram of haplotypes $h_1, h_2$ if it is an ancestor of exactly one of the two haplotypes.

A brick diagram connects a pair of haplotypes via their most recent common ancestors (MRCAs), and it can be visualized as follows.

**Example 1.C. (Example from Figure 1b)** The following is the brick diagram representing the symmetric difference of haplotypes 2 and 3 in Example 1. The symmetric difference between these haplotypes is $\{a, b\}$, and these are the mutations that appear on the brick diagram.

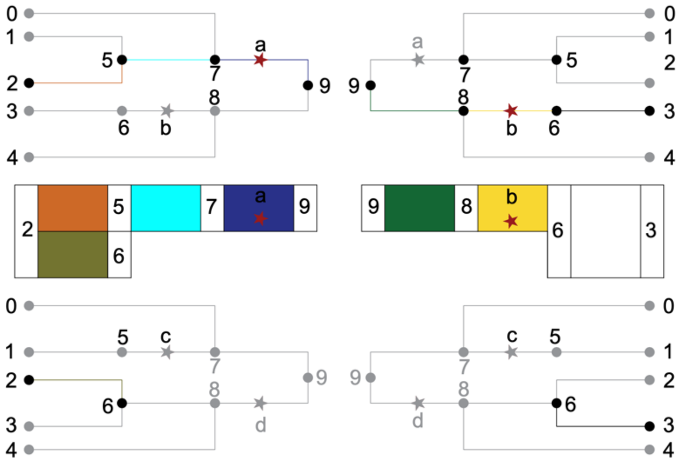


#### 2.3.2 Paths between mutations and definition of the LDGM

The mutations occurring on the bricks of the brick diagram are the symmetric difference between the haplotypes, since bricks have the same descendants at every position (unlike edges of a tree sequence in general). To find the LDGM that satisfies the first condition of Problem 2, we need a set of rules guaranteeing that the labeled bricks of every brick diagram form a connected set in the LDGM.

Consider two labeled bricks $a$ and $b$ in a brick diagram between two haplotypes $h_1$ and $h_2$. Our goal is to create a graph with a path between $a$ and $b$. There are two cases:

First, suppose that $a, b \in h_1$. Both mutations occur on one side of the brick diagram (with respect to the MRCA). There are two subcases: a parent-child relationship if $a$ and $b$ occur on bricks with a shared position, or a recombination (parent-child-parent) relationship if not. In each subcase, we make paths between the bricks as follows:

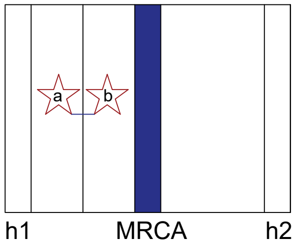


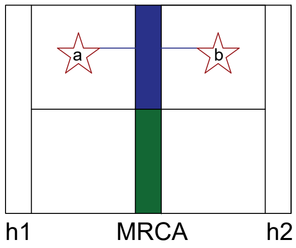


Second, suppose that $a \in h_1, b \in h_2$. The mutations occur on opposite sides of the brick diagram, and we create paths between them that cross through an MRCA. If $a$ and $b$ have overlapping positions, then their MRCA is an MRCA of the haplotypes $h_1$ and $h_2$, and this is a child-parent-child relationship. If not, the path must pass through an MRCA and a child haplotype, and this is a child-parent-child-parent relationship.

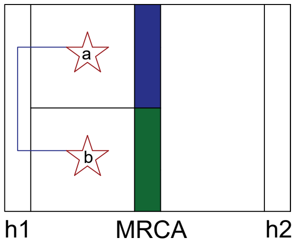


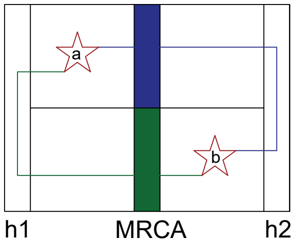

In summary, there are four types of paths in a brick diagram:
1. “I-shaped paths” between parent and child.
2. “V-shaped paths” between two parents of a shared child.
3. “A-shaped paths” between two children of a shared parent.
4. “N-shaped paths” formed by joining an A-shaped and a V-shaped path.
Similar to the recombination-free case (Theorem 1), if any path connecting SNPs $a$ and $b$ contains a third SNP $c$, then $a$ and $b$ are independent given $c$. Therefore, a path between two SNPs (labeled brick) is called *blocked* if it contains another labeled brick. We want the LDGM to have an edge between any two labeled bricks connected with an unblocked path.
**Definition:** The LDGM associated with a bricked tree sequence $T$ is the graph whose vertices are the set of SNPs $S$ and whose edges are as follows:

- If $a$, $b$ are mutations whose bricks have some shared position, then they have an edge in the LDGM if they have an unblocked “I” or “A” shaped path in every tree where they both exist.
- If $a$, $b$ are mutations whose bricks have no shared position, then they have an edge in the LDGM if for some haplotype $h$ in the tree sequence, each of them has an unblocked “V” or “N” shaped path to $h$ at every position where they exist.

**Example 1.D. (Example from Figure 1c)** The following graph is the LDGM of the tree sequence in Example 1.A. For instance, $ab$ is created from an A-shaped path shown in example 1.C.

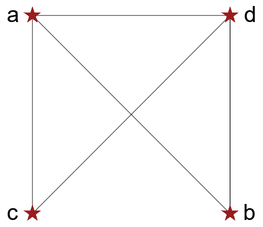


**2.3.3. Correctness of the LDGM**
In this subsection we will show that the LDGM of a tree sequence $T$ is an I-Map for its set of haplotypes $H$.
**Theorem 2.1.** The LDGM is an I-map.
**Proof:** Suppose $h_1, h_2 \in H$ are haplotypes with $a, b \in h_1 \Delta h_2$. We need to show that the LDGM has a path between $a$ and $b$ that stays in $h_1 \Delta h_2$. By the definition of the brick diagram every mutation in $h_1 \Delta h_2$ is contained in the brick diagram $D$ associated with $h_1$, $h_2$. The mutations $a, b$ may have one of our four types of paths. If the path between them is unblocked, then there is an edge between $a, b$ in the LDGM. If a third mutation $c$ blocks the path between $a$ and $b$, it is enough to show that $a$ does not block the path between $c$ and $b$, and $b$ does not block the path between $a$ and $c$ (because there are finitely many mutations). Consider the four types of paths:
**Case 1.** There is an “I” shaped path between an ancestor $a$ and a descendant $b$. If $c$ blocks this path, then $c$ is the descendant of $a$ and the ancestor of $b$, and there are “I” shaped paths between $c$ and both $a$ and $b$.
**Case 2.** There is an “A” shaped path between $a$ and $b$. If $c$ blocks this path, then $c$ is an ancestor of either $a$ or $b$, and hence $a$ does not block the path (“A” shaped path or “I” shaped path respectively) between $c$ and $b$.
**Case 3.** There is a “V” shaped path between $a$ and $b$. Therefore $a$ and $b$ recombined in a haplotype $h$. If $c$ blocks this path, then $c$ is a descendant of $a$ or $b$ and an ancestor of $h$. Therefore, $a$ does not block the path between $c$ and $b$.

**Case 4.** There is an "N" shaped path between $a$ and $b$. If $c$ blocks this path, then based on the position of $c$ on the path between $a$ and $b$ this case is equivalent to either Case 2 or Case 3.

■

#### 2.3.3. Minimality of the LDGM

In this subsection, we will show that the LDGM is minimal in the following sense. For a bricked tree sequence $T$ with mutations $S$, we say that a set of haplotypes $X \subset 2^S$ satisfies the *four gametes condition* with respect to $T$ if $X$ contains the empty haplotype $h_0$ and if for all pairs of SNPs $a, b$:

- $X$ contains no haplotype with $ab$ if and only if $a$ and $b$ are siblings in $T$.
- $X$ contains no haplotype with $aB$ if and only if $a$ is a parent of $b$.
- $X$ contains no haplotype with $Ab$ if and only if $a$ is a child brick of $b$.

**Theorem 2.2.** Suppose $X$ satisfies the four gametes condition with respect to a bricked tree sequence $T$, and let $G$ be the LDGM associated with $T$. For every edge edge $(a, b)$ of $G$, for every third SNP $c$, there exist $h_1$, $h_2$ in $X$ such that $h_1 \Delta\, h_2$ contains $a$ and $b$ but not $c$.

**Proof:** Suppose $a$ and $b$ are adjacent SNPs in the LDGM. There are two cases and several subcases:

**Case 1.** $a$ and $b$ have a shared position. Then they either have a parent-child relationship or, if not, a sibling-sibling relationship.

**Case 1.1.** If $a$ and $b$ have an unblocked "I" shaped path, with $a$ being the ancestor of $b$, consider the child haplotype $h_1$ of $b$. $h_1$ has both $a$ and $b$; suppose it also has a third mutation $c$. Because $c$ does not block the path from $a$ to $b$, either $c$ is an ancestor of $a$, or $c$ is not an ancestor of $b$. If it is an ancestor of $a$, then there exists a haplotype in $h_2 \in X$ with $c$ but not $a, b$, and $h_1 \Delta h_2$ contains $a, b$ but not $c$. Otherwise, if it is not an ancestor of $b$, then some $h_2 \in X$ has derived allele $b$ but not $c$. $h_2$ must also have $a$; the symmetric difference between $h_2$ and the empty haplotype $h_0$ contains $a$ and $b$ but not $c$.

**Case 1.2.** If $a$ and $b$ have an unblocked "A" shaped path, consider the child haplotype $h_1$ of $a$ and the child haplotype $h_2$ of $b$. Suppose that $h_1$ but not $h_2$ has mutation $c$. There are two possibilities, depending on whether $c$ is an ancestor of $a$. Suppose $c$ is not an ancestor of $a$; similar to the parent-child case, $X$ contains a descendent haplotype $h_3$ of $a$ that does not have $c$. The symmetric difference between $h_3$ and $h_2$ contains $a$ and $b$ but not $c$. Now suppose $c$ is not an ancestor of $a$. Because it does not block the path from $a$ to any of the MRCAs of $a$ and $b$, it must be an ancestor of one of those MRCAs. This does not mean that it is an ancestor of $b$, which may not be a descendent of the common ancestor at that particular position (bricks have the same descendant haplotypes at every position, but not the same descendant bricks); in fact, it must not be an ancestor of $b$, since $h_2$ does not have $c$. Therefore, the positions of $c$ and $b$ must not overlap, and in particular, $c$ and $b$ are not siblings. By the four gametes condition, some haplotype $h_3 \in X$ has both $b$ and $c$. This haplotype cannot have $a$, since $a$ and $b$ are siblings; therefore, the symmetric difference between $h_1 \supset (aBc)$ and $h_3 \supset (AbC)$ contains $a$ and $b$ but not $c$.

**Case 2.** Suppose that $a$ and $b$ do not have a shared position. Again, there are two possible relationships that might produce the edge between them: either $a$ and $b$ have a shared descendent $h$, or $a$ has a shared descendent with a sibling of $b$.

**Case 2.1.** Suppose that $a$ and $b$ have an unblocked "V" shaped path. Suppose that $h$ also has some mutation $c$ (otherwise, its symmetric difference with the empty haplotype contains $a$ and $b$ but not $c$).

**Case 2.1.1.** If $c$ is at a different position from both $a$ and $b$, then it is not their ancestor, and by the four gametes condition, there is some haplotype $h_1 \in X$ that has $a$ but not $c$, and there is some haplotype $h_2 \in X$ that has $b$ but not $c$. If $h_1 = h_2$ then the symmetric difference between $h_1$ and the empty haplotype contains $a$ and $b$ but not $c$. If $h_1 \neq h_2$ , then the symmetric difference between $h_1 \supset (aBC)$ and $h_2 \supset (AbC)$ contains $a$ and $b$ but not $c$.

**Case 2.1.2.** Suppose that $c$ has a shared position with $a$. Then $c$ is either an ancestor, a descendent, or a sibling of $a$. We claim that $c$ is an ancestor of $a$. First, $c$ cannot be a sibling of $a$, because sibling bricks never have common descendants. Second $c$ cannot be a descendent of $a$, as any descendent brick of $a$

that is an ancestor of $h$ would block the path between $a$ and $h$. Because $c$ is an ancestor of $a$, $b$ is not an ancestor of $c$, as $b$ is not an ancestor of $a$. By the four gametes condition, there exists some haplotype $h_1 \in X$ that has $c$ but not $b$. If $h_1$ does not have $a$, then the difference between $h$ $(abc)$ and $h_1$ $(ABc)$ contains $a$, $b$ but not $c$. If $h_1$ does have $a$, then let $h_2$ be a haplotype that has $c$ but not $a$. If $h_2$ has $b$, then the symmetric difference between $h_2 \supset (Abc)$ and $h_1 \supset (aBc)$ contains $a$ and $b$ but not $c$. If $h_2$ does not have $b$, then the symmetric difference of $h_2 \supset (ABc)$ and $h \supset (abc)$ contains $a$ and $b$ but not $c$.

**Case 2.2.** Suppose that $a$ and $b$ have an unblocked "N" shaped path, and in particular that $b$ has a shared descendent $h_2$ with some sibling of $a$. Let $h_1$ be the child haplotype of $a$; $h_1$ has $a$ but not $b$. First, suppose that $h_1$ but not $h_2$ has a third mutation $c$. Because $c$ does not block the path from $a$ to $b$, and because it is not an ancestor of $b$, it must not have a shared position with $b$. Therefore, it is not a sibling of $b$, and some haplotype $h_3 \in X$ has both $b$ and $c$. If $h_3$ does not have $a$, then the symmetric difference between $h_3 \supset (Abc)$ and $h_1 \supset (aBc)$ contains $a$ and $b$ but not $c$. If $h_3$ does have $a$, then let $h_4 \in X$ be a haplotype having $c$ but not $a$ (which exists because $c$ is not a descendent of $a$). If $h_4$ does not have $b$, then the symmetric difference between $h_3 \supset (abc)$ and $h_4 \supset (ABc)$ contains $a$ and $b$ but not $c$. If $h_4$ does have $b \supset (Abc)$, then its symmetric difference with $h_1 \supset (aBc)$ contains $a$ and $b$ but not $c$.

■

### 3. Algorithm for LDGM

To build the LDGM, first, we construct the bricked tree sequence. Second, we create two intermediate graphs called the brick-haplotype graph and the SNP-haplotype graph. Third, we remove the haplotypes and construct the LDGM.

#### 3.1. Construction of the bricked tree sequence

Before using a tree sequence to construct an LDGM, we must first modify the tree sequence to create a *bricked tree sequence* using the efficient operations implemented in *tskit*[2–4].

To convert a tree sequence into a bricked tree sequence, we bifurcate an edge to create two bricks whenever recombination changes the edge's descendants between adjacent marginal trees. The tree sequence data structure allows this to be efficiently computed, since adjacent marginal trees in a tree sequence differ by one or more subtree-prune-and-regraft (SPR) operations. Any edges in the subtree which is regrafted in an SPR operation will retain the same descendants at both marginal trees and thus will not need to be bifurcated. Any edges *above* the subtree being regrafted may have different descendants, until the recombination is resolved at the first shared ancestral edge between the origin (at the left marginal tree) and destination (at the right marginal tree) of the subtree being regrafted. Any edges older than the MRCA will have the same descendants in the adjacent marginal trees and thus will not need to be bifurcated.

This algorithm is efficiently implemented using the *tskit.edge_diff()* function, which returns the edges that differ between marginal trees in a tree sequence. The software package *ldgm* efficiently implements this operation using the *tskit* tree sequence development toolkit.

#### 3.2. Construction of the LDGM:

Recall the definition of the LDGM: the graph with SNPs set as the set of vertices and edges as follows.

- If $a$, $b$ are mutations with some shared position, then they have an edge in the LDGM if they have an unblocked path in every tree where they both exist, forming an "I" or "A" shaped path.
- If $a$, $b$ are mutations with no shared position, then they have an edge in the LDGM if for some haplotype h, each of them has an unblocked path to h at every position where each brick exists, forming a "V" or "N" shaped path.

To construct the LDGM from the bricked tree sequence, we start by identifying "I" shaped paths between labeled bricks. We also identify "I" shaped paths between bricks and haplotypes, where recombination might occur. By the way the paths are constructed, we identify which paths are blocked and which ones are unblocked. Second, we identify "A" shaped paths, between pairs of labeled bricks and from labeled bricks to haplotypes, distinguishing blocked and unblocked paths. Third, we identify unblocked "V" shaped paths, which are unions of two unblocked "I" shaped paths, and "N" shaped paths, which are unions of one unblocked "I" and one unblocked "A" shaped path.

Complicating this process is that an edge is only produced between two mutations if they have an unblocked path at *every* position where both bricks exist, not at *any* position. Therefore, it is necessary to keep track of both unblocked and blocked paths, forming edges between mutations only if they have an unblocked path and no blocked path. Tracking of blocked and unblocked paths is done by using "before" and "after" vertices for each brick: before a path goes through a labeled brick, it follows "before" vertices, and afterward, it follows "after" vertices. The "before" and "after" vertices allow paths to "remember" whether they have passed through a labeled brick or not.

Similarly, it is necessary for paths to "remember" whether they are going up the tree or coming down, which is done by using "Up" and "Down" vertices for each brick. The "Up" vertex of a child brick points to the "Up" vertex of its parent brick, and the "Down" vertex of a parent points to the "Down" vertex of its child.

Finally, we use a "U-Turn" vertex to create a separate path for sibling-sibling relationships as a Child-Parent-Child path. This will allow us to choose a stronger edge weight, if possible, for sibling-sibling relationship which preforms better in practice.

The following subsections describe our algorithm in detail.

**3.2.1. Brick-haplotype Graph:**

Our approach to building the LDGM is first to create a graph called a brick-haplotype graph. The brick-haplotype graph contains 6 or 4 vertices for each brick and 2 vertices for each haplotype. For each "labeled" brick with a mutation, we define a cluster of six vertices: Up-Before, Down-Before, Up-After, Down-After, U-Turn, and Out. (The U-turn vertices only affect path weights, not the paths themselves; see Section 3.2.5). The associated cluster with each unlabeled brick has four vertices: Up-Before, Down-Before, Up-After, and Down-After.

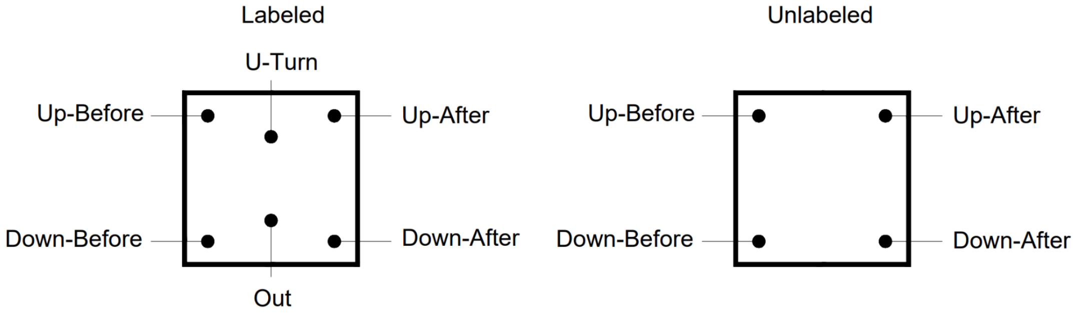


Finally, for each haplotype, we define a cluster of two vertices: Before and After. By assigning multiple vertices to a brick or a haplotype, it effectively "remembers" whether a given path passed through a labeled brick or not. Moreover, considering our four desired types of paths we do not want paths to go down and up within a tree.

The union of these clusters forms the vertex set of the brick-haplotype graph.

The directed edges of the brick-haplotype graph are defined as follows in order to build all "I", "A", "V", and "N" shape paths.

1. "I" shaped paths: Between parent brick (top) and child bricks (bottom) we have the following edges:

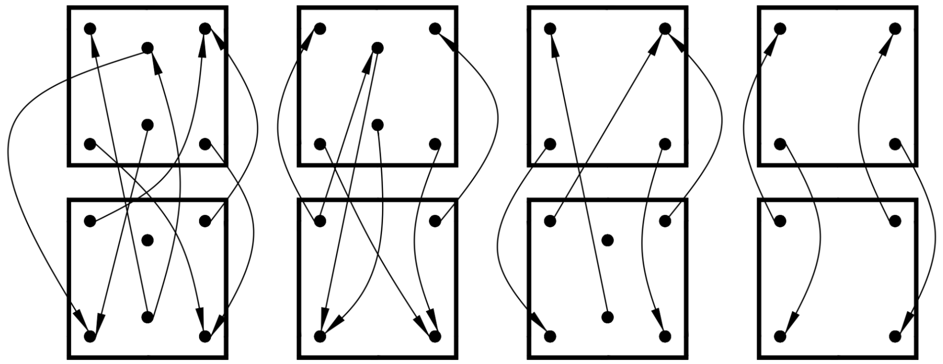

The four subcases are that both bricks are labeled (left), only the parent brick is labeled (left-middle), only the child brick is labeled (right-middle), and neither brick is labeled (right).
The outgoing edges from any labeled brick are:

- Out to Before (Up-Before/Down-Before if the next brick is a parent/child respectively). These edges are the starting edges of paths to create the relationships, and these paths always start at before vertices because they have not been "blocked" by a labeled brick.
- Out of child to U-Turn of labeled parent, and U-Turn of labeled parent to Down-Before of child. The combination of these two types of edges creates sibling-sibling relationships with larger path weights (see section 3.4).
- Down (Before/After) of labeled parent to Down-After of a child. These edges indicate that we have already visited a labeled brick.
- Up (before/after) of labeled child to Up-After of a child. These edges indicate that we have already visited a labeled brick.

From any unlabeled brick, the outgoing edges are:

- Down-Before of an unlabeled parent to Down-Before of the child brick.
- Down-Before of unlabeled parent to Down-Before of the child brick.
- Up-Before of an unlabeled child brick to Up-Before of the parent brick and the U-turn of the parent if it is labeled.
- Up-After of an unlabeled child brick to Up-After of the parent brick.

With these edges, any path stays in the before layer until it visits a labeled brick, then switch to the after layer. Also, any path that is going down does not subsequently go up (it can go up and then down via sibling edges or U-Turn vertices). The U-Turn vertices create an alternative path, potentially with a higher edge weight, between sibling bricks passing through the shared parent (see below).

2. "A" shaped paths: Two bricks are siblings if they have a shared parent brick. Between sibling bricks, we will add the following edges:

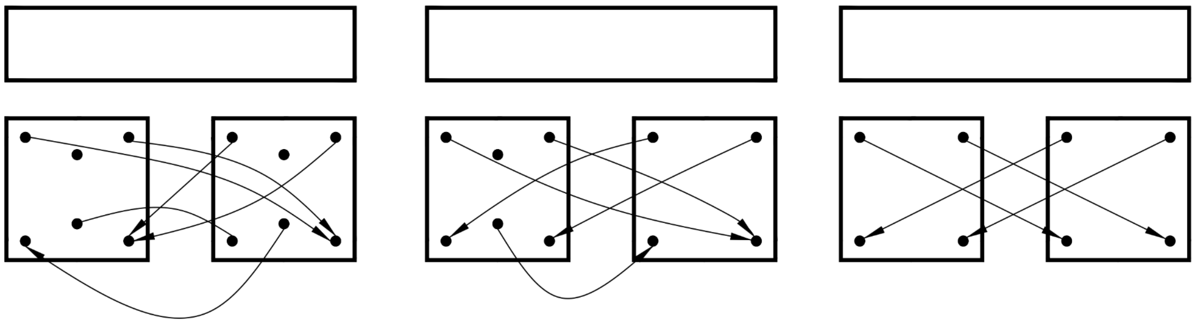

From a labeled brick, the edges are:

- Out to Down-Before. Note that we always start the paths from out vertices; therefore, this type of edges ensures that we have not seen any labeled brick yet.
- Up vertices to Down-After to ensure that the path remembers that it already passed through a labeled brick.

From Unlabeled bricks the edges are:

- Up-Before to Down-Before
- Up-After to Down-After

With these edges we ensure that a path switches to the after level as soon as it visits a labeled brick.

**3.2.2. SNP-haplotype graph:**

The *SNP-haplotype graph* is the reduced graph of brick-haplotype graph such that the vertex set includes one vertex per labeled brick (the set of SNPs) and two vertices per each Haplotype (before and after). We create the SNP-haplotype graph by two-step reductions of the brick-haplotype graph. Note that if a brick has multiple mutations we pick one representative mutation since there is no distinction between mutations on the same brick with respect to conditional dependencies.

**First reduction:** We first create the brick-haplotype graph only using the "I" shaped paths. Then we reduce this graph to produce a "SNP-haplotype graph", as follows. We pick an out vertex of a labeled brick, say $v$, and create the Dijkstra paths starting from the out vertex. For each labeled brick $u$, if one of its before vertices is in the reach set of $v$ but the after vertices are not, then the "I" shaped path between $u$ and $v$ is unblocked, and hence we add an edge between the associated SNPs in the SNP-haplotype graph. Similarly, if the before vertex of a haplotype $h$ is reachable from $v$ but the after vertex is not, then we add a directed edge from the associated SNP to the haplotype. After the reduction, the resulting SNP-haplotype graph contains an edge between two SNPs if there is an unblocked "I" shaped path between them. Similarly, it contains a directed edge from a SNP to a Haplotype if there is an unblocked "I" shaped path from the labeled brick to the Haplotype.

**Second Reduction:** Now, we add the "A" shaped paths to the brick-haplotype graph, reduce the graph to the set of labeled bricks and haplotypes similar to the previous step, and update the edges of the SNP-haplotype graph. The reason for a two-step approach is that when constructing "V" and "N" shaped paths, it is necessary to distinguish between "I" and "A" shaped paths. "V" shaped paths are the union of two "I" shaped paths, and "N" shaped paths are the union of one "I" and one "A" shaped path, but the union of two "A" shaped paths is not an allowed path. The reduction, similar to the previous step, also uses the Dijkstra algorithm. Now two SNPs are adjacent in the SNP-haplotype graph if there is an unblocked "I" shaped or "A" shaped path between them. Note that in the second reduction, we reduced the brick-Haplotype graph only to the set of SNPs; hence, we did not add any new edges to Haplotype vertices. The relationship between bricks (or SNPs) and haplotypes are essential when there is a parent-child relationship since these edges will decode recombinations.

**3.3. Haplotype removal:**

To create the LDGM, we only need to create the edges corresponding to "V" and "N" shaped paths. If two SNPs have edges to a haplotype in the SNP-haplotype graph, then there is a "V" shaped path or an "N" shaped path between them. Therefore, we add an edge between two SNPs in the SNP-haplotype graph if they point to the same Haplotype. Finally, by removing all Haplotype vertices from the SNP-haplotype graph, the remaining graph is the LDGM. The vertices are SNPs, and two SNPs are adjacent if:

1. They have shared positions, and an unblocked path in every tree where they both exist, forming an "I" or "A" shaped path.
2. They have no shared position, and they have an edge in the LDGM if, for some haplotype $h$, each has an unblocked path to $h$ at every position where each brick exists, forming a "V" or "N" shaped path.

**3.4. Weights**

In this subsection, we assign weights to the edges of the brick-haplotype graph using a well-motivated heuristic. LDGMs can have numerous weak edges due to distant genealogical relationships and rare recombination events. We eliminate weak edges by applying an edge-weight threshold, and the choice of threshold allows us to trade accuracy for sparsity.

Edge weights between adjacent bricks are defined by their respective frequencies. Suppose that a parent brick has frequency $f_1$, and a child brick has frequency $f_2$; if these bricks have mutations $a, b$ respectively, then the frequency of observing both mutations is equal to $f_2$. The squared correlation between a mutation on the parent brick and a mutation on the child brick is:

$$r^2_{\text{parent}-\text{child}} = \frac{(f_2 - f_1 f_2)^2}{f_1(1-f_1)f_2(1-f_2)} = \frac{f_2(1-f_1)}{f_1(1-f_2)} = O_2/O_1$$

where $O_a = f_a/(1-f_a)$ denotes the odds of brick $a$. This correlation is small when $f_1 \gg f_2$.

Now, consider a sibling relationship, where the siblings have frequencies $f_1, f_2$. Since siblings have no shared descendants, their squared correlation is:

$$r^2_{sib-sib} = \frac{f_1 f_2}{(1-f_1)(1-f_2)} = O_1 O_2$$

which is small when $f_1 + f_2 \ll 1$. This correlation is one when $f_1 + f_2 = 1$, which occurs when every haplotype is descended from the shared parent of the two siblings, and the expression is greater than one when $f_1 + f_2 > 1$, which does not occur when the siblings have no shared descendants (in these cases, we threshold it at one).

These correlations factor across parent-child relationships. If haplotype 1 is the parent of 2, and 2 the parent of 3, then the squared correlation between 1 and 3 is $r^2_{13} = r^2_{12} r^2_{23}$. The same occurs if 1 and 2 are siblings. This allows us to interpret $d = -\log r^2$ as a distance and to sum distances across paths:

$$d_{12} = -\log r^2_{12}.$$

We observed that sibling-sibling relationships between low-frequency mutations always have small weights but that sometimes, these relationships are important in the LDGM, and excluding them leads to high mean-squared error. This occurs when the two low-frequency siblings have a shared parent mutation that also has a low frequency, such that the two mutations are weakly correlated but strongly correlated *conditional* on that parent. This observation motivates the use of a "U-Turn" vertex at the shared parent creating a different, stronger edge weight between them. If the children are 1,2 and the parent is 3, the U-turn distance is

$$d_{12} = d_{13} + d_{23} = -\log \frac{O_1 O_2}{O_3^2}$$

which is much smaller than $-\log O_1 O_2$ when $O_3$ is small. With this change, the MSE is similar across SNPs at different allele frequencies (Supplementary Figure 7).

For "V" and "N" shaped paths, the path distance is equal to the sum of the path distances of their component paths. The edge distance between a brick and its child haplotype is zero.

#### 3.5. Precision matrix inference

Our precision matrix inference method is based on the graphical lasso (GLASSO)[5], and in particular the DP-GLASSO ("dual-primal GLASSO") method of Mazumder and Hastie[6]. We compute a precision matrix whose nonzero edges correspond to the edges of the LDGM, using an L1 penalty to produce additional sparsity. Then, we remove the L1 penalty but restrict the unpenalized optimization to the edges that were found in the L1-penalized step.

##### 3.5.1 Modified DP-GLASSO

GLASSO and DP-GLASSO maximize the L1-penalized Gaussian likelihood:

$$\hat{P} = \text{argmax}_P \ \frac{1}{2}(\log|P| \ - \ Tr(PR)) - \lambda||P||_1 \tag{1}$$

where $P$ is the estimated precision matrix, $R$ is the sample correlation matrix, $\lambda$ is the regularization parameter, $|\cdot|$ denotes the determinant, and $||\cdot||_1$ denotes the L1 norm. In brief, DP-GLASSO iteratively updates one row/column of $P$ at a time, solving the following L1-penalized regression problem:

$$P_{j\neq i,i} = \text{argmin}_{w\in\mathbb{R}^{m-1}} \frac{1}{2} w^T (P^{-1})_{j\neq i, j\neq i} w - R_{i,j\neq i} w + \lambda||w||_1. \tag{2}$$

(This is L1-penalized ordinary least squares linear regression of the vector $R_{i,j\neq i}$ on the matrix $(P^{-1})_{j\neq i,j\neq i}$). It does so via entry-wise coordinate descend on $w$. DP-GLASSO differs from the original GLASSO method in that it directly estimates the sparse matrix $P$ rather than its inverse; this difference allows us to restrict DP-GLASSO to the edges of the LDGM.

Our precision matrix inference approach differs from DP-GLASSO in three ways. First, we constrain optimization $(1)$ such that $P$ has nonzero entries only for edges of the LDGM, and this constraint means that we must solve a much smaller lasso regression problem in each coordinate descent update (2). The constrained optimization is:

$$\hat{P} = \text{argmax}_P \ \frac{1}{2}(\log|P| \ - \ Tr(PR)) - \lambda||P||_1$$
$$\text{subject to } P_{ij} = 0 \text{ for } (i,j) \notin E.$$

Notice that with the constraint, the objective function only depends on $R_{ij}$ for $(i,j) \in E$.

Second, we include SNP $i$ itself in its own set of neighbors; this differs from (2), where SNP $i$ is excluded. In the corresponding diagonal entry of $R$, which would normally be one, we place a zero. We found that this change improves the convergence rate in practice, and we provide a heuristic justification below. With these two changes, the optimization at each step is:

$$\text{minimize}_{w\in\mathbb{R}^{|N(i)|}} \frac{1}{2} w^T (P^{-1})_{N(i),N(i)} w - (R - I)_{i,N(i)} w + \lambda||w||_1. \tag{3}$$

where $N(i)$ denotes the neighbors of SNP $i$, including $i$ itself. After solving (3), similar to ref. [6], we update the corresponding row and column of $P$:

$$P_{N(i)\backslash i,i} \leftarrow w_{\backslash i}, \qquad P_{i,i} \leftarrow 1 + \hat{R}_{N(j),j} \cdot w.$$

(The formula for the diagonal entry is the same as DP-GLASSO and GLASSO.)

Third, we solve optimization (3) using an iterative proximal gradient method[7], which differs from the element-wise coordinate descent approach of DP-GLASSO and GLASSO. We made this change in order to avoid working with the dense matrix $P^{-1}$.

**3.5.2. Justification for modified coordinate descent approach**

We justify our modified approach heuristically, referring to ref. [6] for a formal discussion. Let $L = I - P$. Multiplying both sides of the equation

$$R = P^{-1}$$

by *L*, it becomes

$$RL = R - I = P^{-1}L$$

and restricting to column *j*,

$$P(R - I)_j = L_j.$$

This equation already resembles an update equation for the $j$th column of $P$. When we restrict the set of edges to those appearing in the LDGM, the log-likelihood (1) depends only on the corresponding elements of $R$. The algorithm can safely ignore non-neighbor entries not only of $P$ but also of $R$. Indeed, the gradient of the log-likelihood with respect to the entries $P_E$ is:

$$\nabla_{P_E} \log \text{likelihood} = (R_E - (P^{-1})_E)$$

such that when the likelihood is maximized, for every SNP *j*,

$$L_{N(j),j} = \left(P|_{N(j)}\right)(R - I)_{N(j),j}.$$

Without an L1 penalty, this equation suggests the following algorithm:

1. Initialize at $P = I$
2. Loop over SNPs *j* one at a time, setting

$$P_{N(j),j} = I_{N(j),j} - \left(P|_{N(j)}\right)(R - I)_{N(j),j}.$$

With an L1 penalty, the unpenalized update equation is modified to produce the SNP-wise optimization problem (3).

**3.5.3. Detailed description of the algorithm**

In detail, the modified coordinate descent outer loop is as follows. We initialize the precision matrix at $P = I$. We loop through the SNPs $j = 1, \dots, M$ five times with an L1 penalty of 0.1, restricting the optimization to the edges of the LDGM that have an edge-weight distance of at most 4 (heuristically equivalent to $r^2 \geq 1/e^4$). At each iteration, we solve optimization (2) (see below for details on the inner loop), we update row/column *j* of the $P$, and we check that $P$ is still positive definite (by attempting to compute its Cholesky factor). Occasionally it is not, indicating that the lasso regression procedure failed to obtain a good solution at the previous step. Typically this is caused by a too-large step size, so when it occurs, we reduce the minimum step size by half and try the previous lasso regression step again; if reducing the minimum step size three times does not fix the problem, we simply use $w^{(0)}$ as the solution (i.e. the non-L1-penalized solution). We have not observed instances where sporadic convergence issues for individual lasso problems lead to low-quality solutions globally, probably because the unpenalized fallback solution is acceptable within the larger optimization routine.

After iterating through every SNP five times, we repeat the procedure without an L1 penalty, restricting to the edges that were found in the previous step (precision matrix entries smaller than 1e-12 are considered non-edges) and initializing at the previous solution. We iterate through every SNP 25 times; each iteration is faster, because the inner loop is no longer required. This approach of removing the L1 penalty after initially using to find a smaller set of edges is designed to approximate the effect of an L0 penalty (which would lead to a nonconvex optimization).

Details of the inner loop are as follows. We initialize

$$w^{(0)} = \left(P|_{N(i)}\right)R_{N(i),i}$$

where

$$P|_A := P_{AA} - P_{AA^c}\left(P_{A^cA^c}\right)^{-1}P_{A^cA}$$

is the Schur complement of $P$ with respect to the SNPs not in $A$. The inverse of $P|_A$ is equal to $(P^{-1})_A$. $w^{(0)}$ is the solution to optimization (3) when $\lambda = 0$. We update $w^{(0)}$ using the proximal gradient approach of ref[7] . Let

$$S_\gamma(x) = \text{sgn}(x)\min(0, \text{abs}(x) - \gamma).$$

This function maps $[-\gamma, \gamma]$ to zero and $\pm(\gamma + |x|)$ to $\pm|x|$. At iteration *k* of the inner loop, we update $w$ with step size $\gamma^{(k)}$ using:

$$w^{(k+1)} = S_{\lambda\gamma^{(k)}}\left(w^{(k)} - \gamma\nabla F\right)$$

where $F(w) = \frac{1}{2}w^T\left(P|_{N(i)}\right)^{-1}w - R^T_{N(i),i}w$. It can be expensive to compute $\left(P|_{N(i)}\right)^{-1}w$ repeatedly (i.e., solving $\left(P|_{N(i)}\right)x = w$ ) using the approach described in Section 5.1.2. This is a sparse system of equations, but its size scales with the number of SNPs in the LD block, which is much larger than the size of $N(i)$. We solve these equations more efficiently by computing the Cholesky factor of $P|_{N(i)}$. We reorder $P$ to put $N(i)$ last, compute the Cholesky factor $A$ of the reordered $P$, and discarding the rows and columns of $A$ besides $N(i)$. (If $N(i)$ were first instead of last, this would be equivalent to the Cholesky of $P_{N(i),N(i)}$, but here it is the Cholesky of $P|_{N(i)}$ instead). This takes roughly as long as solving $\left(P|_{N(i)}\right)x = w$ once. Then, instead of solving $\left(P|_{N(i)}\right)x = w$ at each iteration, we instead solve $AA^Tx = w$, which is fast.

We initialize the step size at $\gamma^{(0)} = 1$, and at each iteration, we check that the update succeeded in reducing the objective function; if the step size is too large, sometimes the iterative procedure will begin to diverge, causing increases in the objective function. When this occurs, we reduce the step size by half and try again, until reaching a minimum step size of 0.1 (by default) at which point we terminate the loop. Otherwise, we terminate the loop after 10 iterations.

**4. Low rank approximations to the local LD matrix**

Low-rank approximations for the local LD matrix have been used by H. Shi et al.[8,9] and others [10,11]. This approach is useful statistically with small reference panels, as small eigenvalues in a reference LD matrix become large and noisy when that matrix is inverted. It is also useful computationally, as the amount of data required to store a rank-*k* LD matrix is similar to the amount required to store a degree-$k$ LDGM precision matrix.

We distinguish between two approximations: the rank-*k* representation of the LD correlation matrix, $R_k = U_k\Lambda_kU_k^T$, and the rank-*k* representation of its inverse, $P_k = U_k\Lambda_k^{-1}U_k^T$. $R_k$ and $P_k$ are not inverses of each other, and their product is the projection matrix $U_kU_k^T$. $R_k$ and $P_k$ have different statistical properties. On the one hand, $R_k$ can be quite a good approximation to $R$, with an MSE similar to that of the LDGM precision matrix inverse for $k$ around 50 (see Supplementary Figure 4). It makes sense that $R_k$ works well by this metric: the MSE is

$$MSE_k = \frac{1}{m^2}\sum_{j=k+1,\ldots,m}\lambda_j^2$$

the sum of the squares of the small eigenvalues. On the other hand, $P_k$ is not an especially good approximation to $R^{-1}$. $P_kR$ is the projection matrix onto the span of $U_k$, which means that $P_kR^2 = R_k$ is expected to be a good approximation for $R$, but $P_kR$ is a poor approximation for $I$.

To quantify this, we define the *alternative MSE ratio* as:

$$f(P) := \frac{Tr\big((I - PR)(I - RP)\big)}{Tr(PRRP)}.$$

This ratio quantifies how close $PR$ and $RP$ are to $I$, while guaranteeing that when $P$ is nearly the all-zeros matrix, the error is large. The numerator quantifies the mean-squared difference between a random vector $x$ and the vector $PRx$. If $x \sim N(0, I)$:

$$E\left(||x - PRx||^2\right) = Tr\left((I - PR)^T(I - PR)E(xx^T)\right)$$
$$= Tr\left((I - RP)(I - PR)\right).$$

Supplementary Figure 4 shows the alternative MSE ratio for the LDGM precision matrix compared with $P_k$ for different values of $k$. Even with $k = 500$, the alternative MSE ratio is much higher for $P_k$ than for the LDGM precision matrix.

These error metrics imply that a low-rank approximation is appropriate for some applications, but not for others. In applications that involve multiplying a vector by $R_k$ (e.g., simulating GWAS summary statistics), the approximation is satisfactory. In applications that involve $P_k$, which seems to include existing methods, we caution that large $k$ might be required to obtain adequate accuracy. The best choice of *k* might depend on whether within-sample or out-of-sample LD is used, with better within-sample performance at large *k* and better out-of-sample performance at small *k*.

### 5. Calculations involving the LDGM precision matrix

In this section, we describe how LDGM precision matrices can be used to perform useful computations. Section 5.1 describes basic operations, and section 5.2 describes calculations involving a Gaussian model for the effect size distribution (i.e., the model underlying BLUP).

#### 5.1 Basic matrix operations

When deriving efficient formulas involving LDGM precision matrices and GWAS summary statistics, an important consideration is that most of the time, some of the SNPs in the LDGM will be missing from the GWAS summary statistics. (It is normally less significant if SNPs in the summary statistics are missing from the LDGM; see below). Missingness cannot be handled by discarding the rows and columns of the precision matrix that are missing in the summary statistics, since the resulting matrix is not the precision matrix of the non-missing SNPs. Let $R$ be a correlation matrix, let $P = R^{-1}$ be a precision matrix, and let the indices 0,1 denote missing and non-missing SNPs respectively, such that

$$R = \begin{pmatrix} R_{00} & R_{01} \\ R_{10} & R_{10} \end{pmatrix}, \qquad P = \begin{pmatrix} P_{00} & P_{01} \\ P_{10} & P_{10} \end{pmatrix}.$$

$R_{11}$is not equal to $P_{11}^{-1}$; instead, it is equal to the Schur complement of $P_{11}$:

$$R_{11}^{-1} = P/P_{00} := P_{11} - P_{10}P_{00}^{-1}P_{01}.$$

In general, when performing precision matrix operations with missing data, there are two choices for how to proceed: either compute the Schur complement explicitly, or perform computations using the definition of the Schur complement that avoid computing it. When a large number of SNPs are non-missing, it is better to avoid computing the Schur complement (which might be dense), but if there are only a handful of non-missing SNPs (which is the case, e.g., in certain computations that arise when computing the precision matrix), it is better to simply compute the Schur complement (in particular, its Cholesky decomposition).

##### 5.1.1 Precision matrix by vector multiplication

Suppose we wish to solve

$$R_{11}x = y \tag{4}$$

given a vector or tall matrix $y$. We do so using the definition of $P/P_{00}$:

$$x = P/P_{00}y = P_{11}y - P_{10}z, \qquad P_{00}z = P_{01}y.$$

This approach is extremely fast compared with directly solving the dense system of equations (4). It is implemented in the *precisionMultiply* function (see Code Availability).

##### 5.1.2. Precision matrix by vector division

Suppose we wish to compute

$$y_1 = R_{11}x_1 \tag{5}$$

given a vector or tall matrix $x$. We do so by filling in zeros for the missing SNPs to form $x = [0\ x_1]^T$ and solving

$$Py = x, \qquad y = [y_0\ y_1]^T$$

and discarding the unwanted entries $y_0$.

Typically, this approach is only modestly faster than evaluating equation (5), depending on the number of SNPs in an LD block (with a larger improvement when there are more SNPs). It is implemented in the *precisionDivide* function (see Code Availability).

##### 5.1.3. Computing the determinant

Suppose we wish to compute the determinant of $R_{11}$, denoted $|R_{11}|$. We do so with the identity

$$|R_{11}| = |P/P_{00}|^{-1} = \frac{|P_{00}|}{|P|}.$$

To evaluate the determinant in a numerically stable manner, we actually compute the log-determinant, using the identity

$$\log|P| = 2\sum_j \log A_{jj}\,, \qquad P = A^T A$$

where $A$ is the Cholesky factor of $P$. Typically, this approach is substantially faster than a similar approach involving $R$.

More generally, we might wish to compute the determinant of a slightly different matrix:

$$|R_{11}^{-1} + T_{11}|$$

where $T_{11}$ is a sparse matrix (for example, see below). To do so, we fill in zeros where there are missing SNPs to form a matrix $T$ of size equal to that of $P$. The Schur complement of $P + T$ is

$$(P+T)/(P+T)_{00} = P_{11} + T_{11} - P_{10}P_{00}^{-1}P_{01} = R_{11}^{-1} + T_{11}$$

so the determinant can be computed as

$$\left|R_{11}^{-1} + T_{11}\right| = \frac{|P+T|}{|P_{00}|}.$$

##### 5.1.4. Computing the likelihood of the GWAS summary statistics under the null

For a null trait ($\beta = 0$), the GWAS *Z* scores, $z$, follow the following distribution[12, 13]:

$$z \sim N(0, R_{11}) \tag{6}$$

Equivalently,

$$(P/P_{00})z \sim N(0, P/P_{00}).$$

The likelihood of (6) is:

$$\log f(z) = -\frac{1}{2}(m \log 2\pi + |P/P_{00}| + z^T(P/P_{00})z).$$

#### 5.2. Calculating the BLUP and related quantities

Let $\beta$ be the vector of causal per-allele effect sizes for a trait $y$, such that

$$y = X\beta + \epsilon$$

where $X$ is a row vector of genotypes and $\epsilon$ is a noise term. The best linear unbiased predictor (BLUP) is the posterior mean of $\beta$ under a Gaussian model:

$$\beta \sim N(0, \Sigma).$$

This model includes as an important special case the infinitesimal model, $\Sigma \propto I$, and it can be generalized by making $\Sigma$ itself a random variable. Moreover, the diagonal entries of $\Sigma$ can depend on the allele frequencies in a manner consistent with known allele frequency-dependent architecture.

#### 5.2.1. *BLUP-ldgm*

The distribution of the GWAS Z scores, $z$, given $\beta$ is:

$$z|\beta \sim N\left(n^{\frac{1}{2}} R D \beta, R\right)$$

where $D$ is a diagonal matrix that converts $\beta$ into normalized units: under Hardy-Weinberg equilibrium,

$$D_{jj} = \left(2p_j(1-p_j)\right)^{\frac{1}{2}}.$$

The unbiased effect size estimates are:

$$\hat{\beta} = n^{-\frac{1}{2}} D^{-1} P z$$

$$\hat{\beta}|\beta \sim N(\beta, n^{-1} D^{-1} P D^{-1}).$$

(Note that these estimates are extremely noisy; the diagonal elements of $P$ can be greater than 10, and the diagonal elements of $D^{-1}$ are also large for low-frequency SNPs). The posterior mean is:

$$\hat{\beta}_{\mathrm{BLUP}} \coloneqq E(\beta|z) = (\Sigma^{-1} + V^{-1})^{-1} V^{-1} \hat{\beta}, \qquad V = n^{-1} D^{-1} P D^{-1}.$$

This simplifies to:

$$\hat{\beta}_{\mathrm{BLUP}} = n^{1/2} \Sigma D (n D \Sigma D + P)^{-1} P z$$

which can be evaluated efficiently as described in sections 5.1.1 and 5.1.2, including when data are missing. This is implemented in the function *BLUPx*.

#### 5.2.2 Simulating summary statistics

LDGMs can be used to simulate summary statistics from their asymptotic sampling distribution (equation 6). First, we choose an effect size covariance matrix $\Sigma$. In our single population simulations, this is chosen to be proportional to $I$, with a scalar that matches the chosen heritability:

$$h^2 = E(\beta^T D R D \beta) = Tr(DRD\Sigma)$$

$$= \sigma^2 Tr(D^2) \text{ when } \Sigma = \sigma^2 I.$$

In the multiple population setting, we concatenate the vectors of effect sizes for each population, and we choose a populations-by-populations covariance matrix, $\Sigma_{\mathrm{pop}}$. The diagonal entries are chosen to match the heritability in each population, and the off-diagonal entries are chosen to match the desired cross-population genetic correlation, i.e.:

$$r_{pop}(a,b) = \frac{\Sigma_{pop}(a,b)}{\sqrt{\Sigma_{pop}(a,a)\Sigma_{pop}(b,b)}}.$$

Then the covariance matrix of the concatenated effect sizes is:

$$\Sigma = I \otimes \Sigma_{\mathrm{pop}}$$

where $I$ is the identity matrix of size equal to the number of SNPs and $\otimes$ denotes the Kronecker product. If any SNPs are missing in some of the populations, the corresponding rows and columns are dropped from $\Sigma$. More generally, the cross-population effect size distribution of each SNP can be chosen from a mixture of normal distributions.

To sample the summary statistics efficiently, first we sample the true effect sizes $\beta$, and then (for each population separately) we compute the mean of their summary statistics as:

$$E(z|\beta) = n^{1/2} P^{-1} D \beta$$

We produce noise with covariance matrix $R$ by computing the Cholesky of $P$, $P = AA^T$, and solving

$$A\epsilon = x, \qquad x \sim N(0, I).$$

The summary statistics are $z = n^{1/2}P^{-1}D\beta + \epsilon$. These equations are implemented in the function *simulate_sumstats_populations.*

##### 5.2.3. BLUP likelihood

The likelihood of the summary statistics under the BLUP model, not given $\beta$, is:

$$z \sim N(0, R + nRD\Sigma DR)$$

where $RD\Sigma DR$ is the covariance matrix of the per-s.d. marginal effect sizes, $\alpha = corr(X, y) = RD\beta$. Equivalently,

$$Pz \sim N(0, P + nD\Sigma D) \qquad (7)$$

The likelihood of equation (7) is:

$$\log f(z) = -\frac{1}{2}(m \log 2\pi + |P + nD\Sigma D| + (Pz)^T(P + nD\Sigma D)Pz) \qquad (8)$$

When there are missing SNPs such that we only observe $z_1$, we assume that their effect sizes are zero. Let $S = D\Sigma D$ and $S_{11}$ denote the non-missing block. Let $x = (P/P_{00})z_1$. Equation (8) becomes:

$$\log f(z_1) = -\frac{1}{2}(m \log 2\pi + |P/P_{00} + nS_{11}| + x^T(P/P_{00} + nS_{11})x)$$

which can be evaluated efficiently as described in sections 5.1.1, 5.1.2, 5.1.3. This is implemented in the function *GWASLikelihood*.

The gradient of the likelihood can also be computed efficiently. Let $y = (D\Sigma D + n^{-1}P/P_{00})^{-1}x$. The partial derivative with respect to $\Sigma_{jj}$ is:

$$\frac{\partial}{\partial \Sigma_{jj}} \log P(z|D) = D_{jj}^{-2}\left(\begin{bmatrix} D\Sigma D & 0 \\ 0 & 0 \end{bmatrix} + n^{-1}P\right)^{-1}_{jj} + D_{jj}^{-2}y_j^2.$$

The first term appears like it would be slow to evaluate, as it is an element of the inverse of a large sparse matrix, and inverting this matrix directly would be slow. However, efficient approaches can be used to extract the diagonal elements of the inverse without computing the entire matrix []. This approach is implemented in the function *GWASLikelihoodGradient*, which computes the derivatives with respect to all of the diagonal elements of $\Sigma$.
